# Searching for influencers among placental immune cells in preeclampsia

**DOI:** 10.1101/2025.03.04.641363

**Authors:** Hyojung Paik, Tae Lyun Ko, Myungsun Park, Jong-Eun Park, Danial Bunis, Marina Sirota, Byung Soo Lee, Hyeong-Sam Heo, Sung Ki Lee

**Affiliations:** Center for Biomedical Computing, Korea Institute of Science and Technology Information (KISTI), and Department of Data and HPC Science, Korea National University of Science and Technology (UST), Daejeon 34141, Republic of Korea; Graduate School of Medical Science and Engineering, KAIST, Daejeon 34141, Republic of Korea; Department of Obstetrics and Gynecology, College of Medicine, Myunggok Medical Research Institute, Konyang University, Daejeon 158, Republic of Korea; CoLabs, Bakar ImmunoX Initiative, University of California, San Francisco, CA94143, USA; Bakar Computational Health Sciences Institute (BCHSI), University of California, San Francisco, CA94158 USA; New Drug Development Center, OSONG Medical Innovation Foundation, Chungbuk 28515, Republic of Korea

**Author notes:** These authors are equally contributed (Hyojung Paik, Tae Lyun Ko).

## Abstract

Cells in maternal and fetal immune systems may communicate, leading to immune tolerance during pregnancy; however, this hypothesis remains controversial. Here, we profiled single-cell transcriptional signatures in placental layers comprising the maternal–fetal interface and deep placenta, then searched for genes associated with preeclampsia. To investigate the underlying principle of the failure of immune tolerance, we started by clarifying the systemic framework, comprising models of immune interaction frequency (IIF) and specific triggers (i.e., influencers) of tolerance (IT). We generated single-cell transcriptional profiles of normal term (Norms) and preeclampsia preterm (PePT) parturitions. Fetal and maternal cells are admixed across the placenta, for both Norms and PePTs, rejecting the IIF model of immune failure during pregnancy posed by excessive interactions between fetomaternal cells. Whereas placental layers are well mixed with maternal cells, we identified a conserved gradual immune transition of fetal T-cells in both PePT and Norm, disproving the IIF model. To search for influencers of PePT in the IT model, we established and validated a classification model for PePT and Norm immune cells, including T-cells, and then prioritized major contributors to the classifier model, which are highly enriched in ligands and receptors (*p* = 5.98e^−5^). Among the prioritized ligand receptors, SPP1 and CD44 are suggested as influencers of inflammation signatures and were experimentally validated by the exclusive colocalization of SPP1- and CD44-expressing cells in the PePT placentas. Different interleukin-4 and interferon-γ levels in the serum and urine of PePTs further support the contribution of SPP1 to associated pathways, including allograft rejection. Our findings provide insight into the influence of specific immune interactions between cells in the human placenta and their influencer-derived impact on PePT.

## Introduction

Allogeneic individuals coexist during pregnancy in mammals, including humans (*1*). The human placenta is a heterogeneous organ critical for establishing the fetomaternal interface and acquiring immunological tolerance between semiallogenic cells. Placental dysfunction, such as the failure of immune tolerance, contributes to significant complications, such as preeclampsia, a potentially lethal hypertensive disorder during pregnancy (*2*). Although preeclampsia has a broad spectrum, severe preeclampsia frequently leads to preterm parturition before 37 completed weeks of gestation, which may contribute to neonatal mortality (*3*). The biological mechanisms that lead to preterm birth preeclampsia and the factors that confer preeclampsia preterm parturitions (PePT) remain inconclusive.

Preeclampsia (PE) is a syndrome of multiple etiologies, characterized by the new onset of high blood pressure (systolic blood pressure ≥140 mmHg and/or diastolic blood pressure ≥90 mmHg) during pregnancy, followed by fatal outcomes for both the mother and fetus (*4*). Thus, the clinical manifestation of preeclampsia implies abnormal angiogenesis during early placentation (*5*). For example, the levels of sFlt-1, an anti-angiogenic protein that antagonizes VEGF (vascular growth factor) and PIGF (placental growth factor) are as higher in women with preeclampsia than in those with normal pregnancies (*4*). This dysfunction of angiogenesis leads to poor structural development of placenta, resulting in hypoxia and oxidative stress and explaning the hypertensive symptoms of preeclampsia.

Alongside the angiogenesis-based hypothesis, an insufficiency in gestational immune tolerance has also been suggested as a pivotal etiology of preeclampsia. During pregnancy, neutrophils, monocytes, and natural killer (NK) cells initiate inflammation, which induces endothelial dysfunction, and activated T-cells may result in inadequate tolerance (*6*). Based on the evidence from single-cell RNA sequencing and uterine organoids, Li et al reported that uterus NK cells affect diverse genetic programs, including regulating blood flow and adequate immune responses contributing to preeclampsia (*7*). Successful pregnancies depend on the delicate interplay between local immune recognition and immune tolerance mediated by NK and T-cells. However, this well-regulated immune interaction may be disrupted in pregnancy-related disease, including preeclampsia (*8*). The exact role of immune cells in reproductive success in humans remains controversial.

The use of new research technologies, including scRNA-sequencing (scRNA-seq,) successfully resolves several immunological questions by deciphering transcriptional signatures consisting of tens of thousands of cells from given tissues (*9*). Roser et al. used single-cell transcriptomics to comprehensively resolve the cellular and molecular states involved in maternal– fetal communication in the decidua during the early trimester (*10*). Focusing on pregnancy outcomes, a previous attempt provided a heterogeneous cellular catalog and transcriptional profiles for normal and sporadic preterm births (*11*). Integrative analysis of cell-free plasma RNA and scRNA-seq data from diverse placentas has revealed promising approaches for understanding the cellular dynamics of pregnancy outcomes (*12*). However, the understanding of well-orchestrated interacionts between heterogeneous immune cells in complex placental layers for successful pregnancy remains incomplete.

Herein, we utilized scRNA-seq to generate a single-cell landscape of the human placenta to compare normal term (Norm, approximately 39 or 40 weeks) and preeclampsia preterm parturitions (PePT, 30.8±2.7 weeks) by examining two placental layers, maternal-fetal interface (MF) and deep-placenta (Pla). All women underwent delivery without labor, and all preterm births had server preeclampsia conditions. We began with a hypothetical framework in which immunological abnormality underpine the etiological mechanisms of PePT. More directly, we investigated the immunological dysfunction (i.e., failure of normal fetomaternal tolerance) of PePT with respect to inflammation resulting from either (i) an immune interaction frequency (IIF) model for excessive semiallogeneic cell interactions, or (ii) an influencer tolerance (IT) model for preeclampsia-specific cellular interactions. We described the cellular composition of each placental layer and then established a supervised classifier to prioritize PePT-associated transcriptional signatures among given cell types, such as T-cells. Interestingly, ligand and receptor genes that handle crosstalk between cells are highly enriched in those of encoding PePT-ssociated transcripts, supporting the existance of *influencers* triggering immunological cascades. Cell-cell interaction analyses were used to infer PePT-specific communication networks among cells in each tissue, and the results were then validated experimentally using confocal microscopy. Using the Olink profiling, we leveraged existing urine and serum samples to identify associated proteomic signatures, which may be associated with PePT-specific transcriptional impact.

## Results

### A single-cell portrait through the human placenta by pregnancy outcomes

We succinctly investigated the breakdown of immune tolerance through a theoretical divide-and-conquer strategy: one focused on excessive allogeneic cell interactions, whereas the other examined immune harmonization failure triggered by specific factors. Figure 1 depicts our bifocal approach and study design. As shown in Fig. 1A, the IIF model suggests that excessive semiallogeneic cell interactions would mainly lead to a state of abnormal inflammation in pregnant women, resulting in PePTs, as suggested by the finding that in NK-cell associated poor-angiogenesis in PePTs contributes to hypoxia and inflammation (*13*, *14*). The IT model suggests that a specific cell-to-cell interaction disrupts the immune tolerance of pregnancy; that is the “Tail wagging the dog”. The premise of the IIF model is the differential dispersion of semiallogeneic cells, such as fetal cells within the maternal placenta, between the deep placenta versus outer layers. Thus, we gathered cells throughout the placental layers, including at the maternal-Fetal interface (MF) and deep placenta (Pla) layers by pregnancy outcomes (i.e., Norm and PePT), and then determined the density of fetal and maternal cells in the corresponding regions (Fig. 1B and C).

**Fig. 1.**
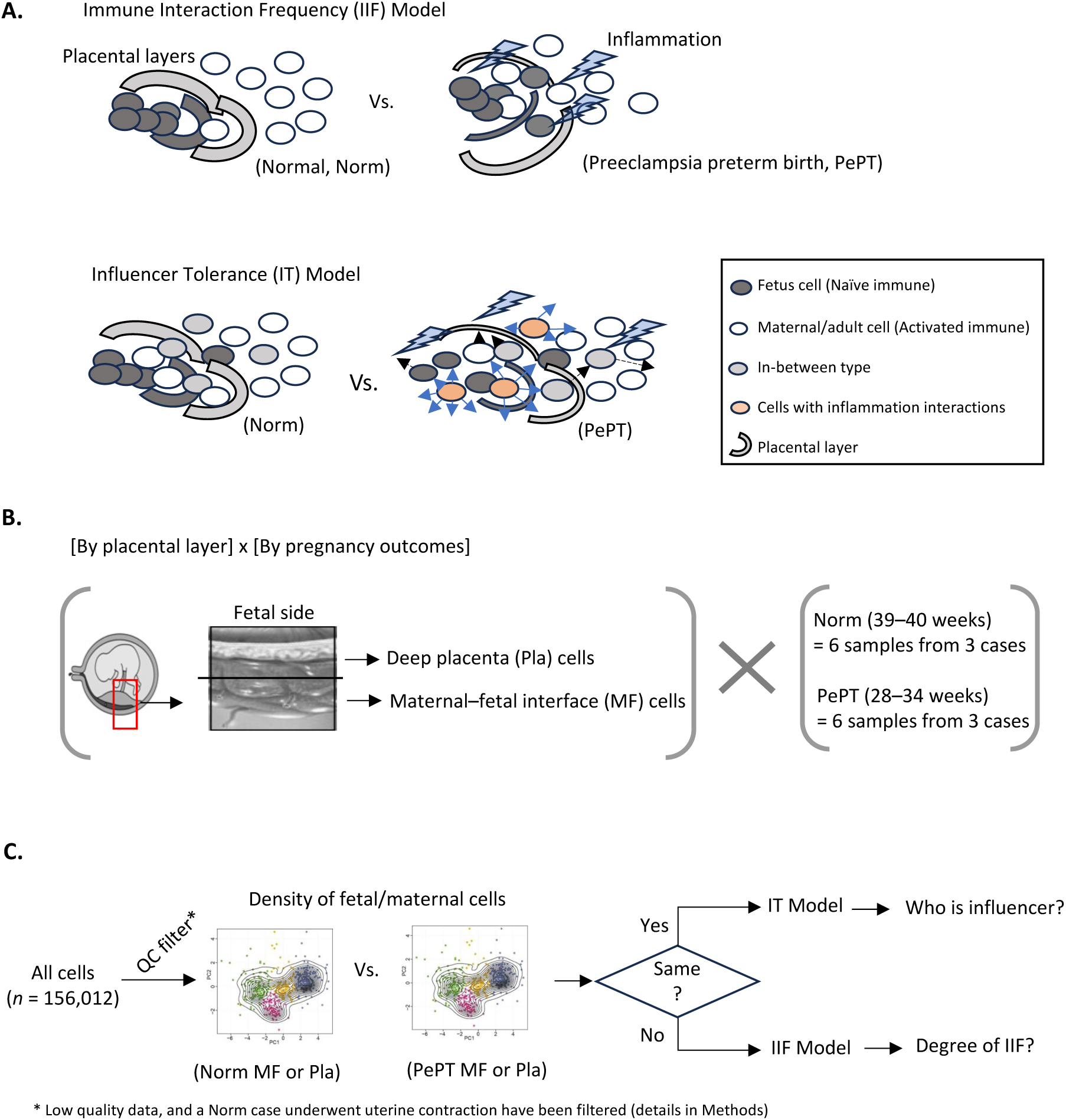
Theoretical and experimental overview. (A) Theoretical models for how immune cell fetal-to-adult harmonization may occur. Upper panel: *immune interaction frequency* (IIF) model, in which a wave of inflammation emanates from the general interactions between nonself and self-cells across placental layers. Lower panel: the *influencer tolerance* (IT) model, in which an original population of fetal cells undergoes multiple, gradual, relatively homogeneous changes that ultimately result in the immune tolerance of normal pregnancy. However, idiopathic interactions can have an adverse effect, causing abnormal inflammation and leading to failure in immune harmonization. (B) An experimental overview shows the tissues and compares pregnancy outcomes to validate our hypothesis. To examine our models, we gathered cells from placental layers, including the maternal–fetal interface (MF) and deep placenta (Pla) layers for each case comprising normal and preeclampsia preterm births. (C) Analysis overview presenting the validation process of the suggested models.

To explore the cellular interactions and distributions between semiallogenic cells in placenta, we first established a molecular portrait of placenta cells from Pla and MF tissues the women who underwent PePT (*n* of individual = 4, *n* of tissues = 8, mean of gestation = 30.8±2.7 week), or who delivered a fetus at full term, Norm (*n* of individual = 4, *n* of tissues =8) (Table 1). In the present study, Pla represents a placental region closer to the fetal side, whereas MF represents the outer layers (Fig. 1B). All 16 tissues were harvested immediately after cesarean delivery, and paired region of Pla and MF were included for each case. For overall recapitulation of the placental layers (i.e., fetal or maternal side), each Pla and MF comprised pooled cells from multiple corresponding regions. The Pla and MF samples from Norm and PePT women were used to prepare for single-cell dissociation followed by scRNA-seq (details in the Materials and Methods section). We used a dual-cross confirmation approach using maternal and fetal genotyping to assign cell type origin during single-cell analysis. First, we clustered cells based on allele variances using Souporcell (*15*) and then reconfirmed the clusters based on the assayed genotypes of peripheral blood monoclear cells (PBMCs) of the mother or cord blood of the fetus (see the Materials and Methods section). Over 1000 alleles (mean of 1115.875± 381.2 per sample) were examined to confirm the inferred origin (i.e., fetal or mother) of cells cells for each sample. Among the 16 tissues collected from eight patients, which consisted of 177,884 cells (Table 1), we omitted two tissues from one individual owing to the poor data quality. Among the twelve tissues (two paired tissue from each patient) from six individuals (three Norms and three PePTs), which indluced more than 130,000 cells, 103,394 cells passed our quality control threshold. We subsequently conducted preprocessing procedure described in the Materials and Methods section. In addition, two paired tissues (i.e., Pla and MF) from one patient comprising over 20,000 cells were separately used for examine the reproducibility of the main analysis.

**Table 1.** Patient statistics.

| Patient ID | Sample ID | Tissue | Gestational week<br>(mean of PePT<br>$30.8 \pm 2.7$) | Outcome<br>(Offspring sex) | Cells | Usage<br>(labor Y/N) |
| --- | --- | --- | --- | --- | --- | --- |
| Norm1 | NormMF1 | MF<sup>1</sup> | 39 + 6 | Norm<sup>3</sup> (female) | 17,648 | QC fail<sup>5</sup> (labor Y) |
|  | NormPla1 | Pla<sup>2</sup> | 39 + 6 | Norm (female) | 7464 |  |
| PePT2 | PePTMF2 | MF | 32 + 1 | PePT<sup>4</sup> (female) | 8771 | Main analysis<br>(labor N) |
|  | PePTPla2 | Pla | 32 + 1 | PePT (female) | 4571 |  |
| PePT3 | PePTMF3 | MF | 28 + 6 | PePT (male) | 12,262 |  |
|  | PePTPla3 | Pla | 28 + 6 | PePT (male) | 11,433 |  |
| Norm4 | NormMF4 | MF | 39 + 1 | Norm (male) | 13,204 |  |
|  | NormPla4 | Pla | 39 + 1 | Norm (male) | 9757 |  |
| PePT5 | PePTMF5 | MF | 34 + 1 | PePT (male) | 15,117 |  |
|  | PePTPla5 | Pla | 34 + 1 | PePT (male) | 8240 |  |
| Norm6 | NormMF6 | MF | 38 + 2 | Norm (male) | 9891 |  |
|  | NormPla6 | Pla | 38 + 2 | Norm (male) | 12,976 |  |
| Norm7 | NormMF7 | MF | 38 + 4 | Norm (female) | 10,923 |  |
|  | NormPla7 | Pla | 38 + 4 | Norm (female) | 13,754 |  |
| PePT8 | PePTMF8 | MF | 28 + 5 | PePT (male) | 10,947 | Validation<br>(labor N) |
|  | PePTPla8 | Pla | 28 + 5 | PePT (male) | 10,926 |  |

<sup>1</sup>MF, maternal–fetal interface of placenta; <sup>2</sup>Pla, deep placenta; <sup>3</sup>Norm, normal full-term birth; <sup>4</sup>PePT, preeclampsia preterm birth, gestational week $\leq 34$ ; <sup>5</sup>Failure of data quality control(QC) threshold including labor of delivery or uterine contraction and lack of expressed genes per cell.

Unsupervised uniform manifold approximation and projection (UMAP) clustering analysis of of those 103,394 cells and cell type annotation identified 24 distinct cell type clusters corresponding to immune and nonimmune cells across the Pla and MF (table S1). Nonimmune cells included stromal cells (two clusters), decidual cells, syncytiotrophoblasts (STB), cytotrophoblasts (CTB), extravillous trophoblasts (EVT), nonproliferative interstitial CTB <npiCTB), endothelial cells, and endometrial cells (Fig. 2A). The immune cell types identified were monocytes, macrophages, NK cells, B-cells and T-cells. The decidual cells (green in Fig. 2A), derived from the maternal uterus (*16*), were identified as maternal cells, and this identity was subsequently confirmed based on presented genotype data from the maternal PBMCs using our dual-confirmation approach. Likewise, the identified origin of the cells provided favorable support for the known origin of cell types, such as EVT cells consisting of fetal cells, allowing us confidence in our analysis (Figure 2B). Based on the placental layers of the corresponding origin of the cells (upper parts of Fig. 2A and 2B), fetal and maternal T/NK cells were interspersed across the placental layers regardless of whether they were Norm or PePTs.

**Fig. 2.**
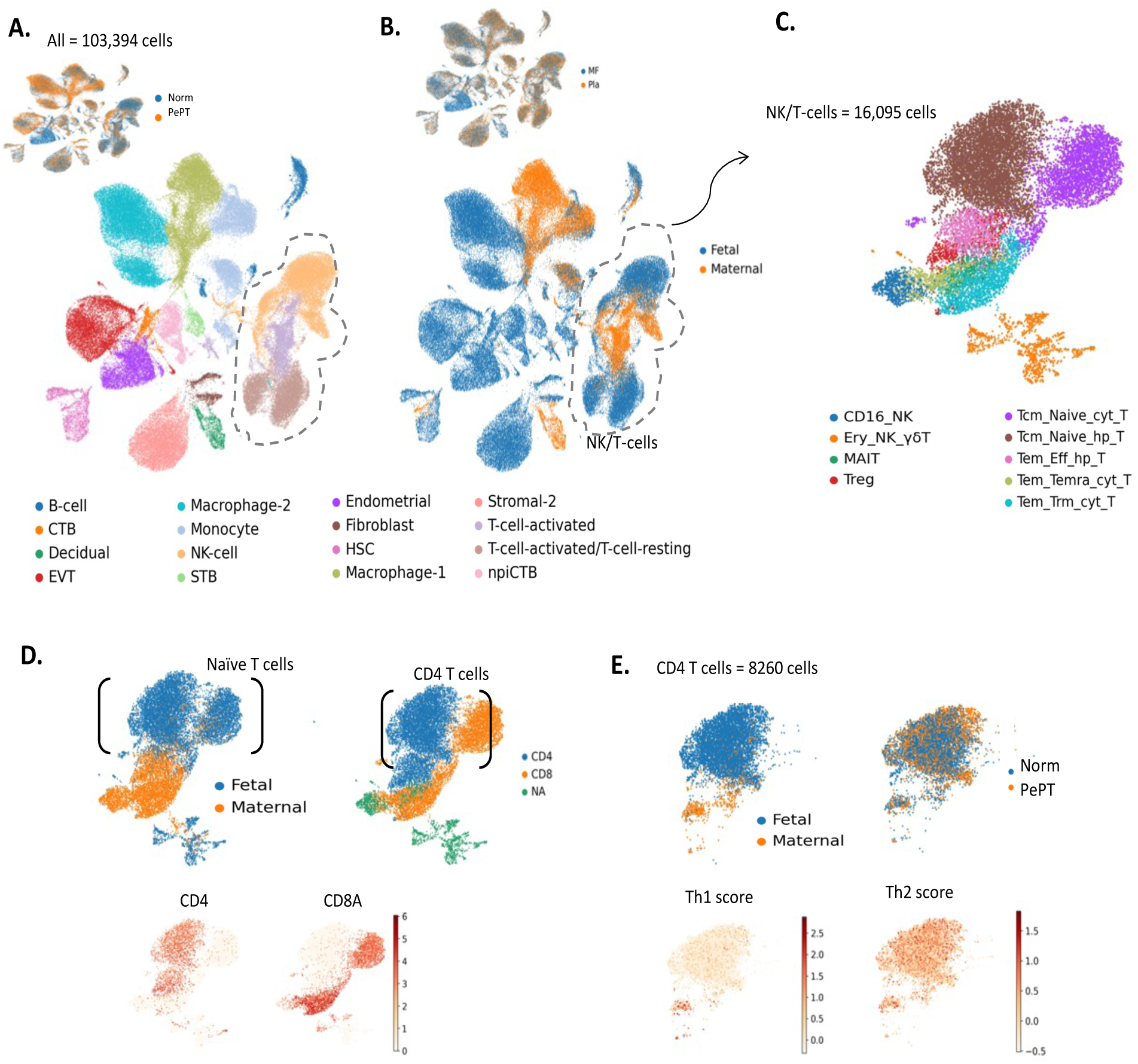
Single-cell-level overview by placental layers, cell origins, and pregnancy outcomes. **(A)** Uniform manifold approximation plot (UMAP) using cells from our study, where dots represent single cells and cell types by color. Endometrial, endometrial-like; STB, syncytiotrophoblast; EVT, Extravillous trophoblast; CTB, cytotrophoblast; HSC, hematopoietic stem cell; npiCTB, nonproliferative interstitial cytotrophoblast. (**B**) Distributions of single cells colored by maternal or fetal origin. (**C**) UMAP using selected T and NK cells from A. MAIT, mucosal-associated invariant T-cell; Treg, regulatory T-cell; Tcm_Naive_cyt_T, T central memory(Tcm) and naïve cytotoxic T-cell; Tcm_Naive_hp_T, Tcm naïve helper T-cell; Tem_Eff_hp_T, T effector memory (Tem) and effector helper T-cell; Tem_Temra_cyt_T, Tem/Temra (effector memory cells re-expressing CD45RA) cytotoxic T-cells; Tem_Trm_cyt_T, Tem/Trm (Tissue-resident memory) cytotoxic T-cells; Ery_NK_γ8T, Erythroid, NK, and γ8T-cells. (**D**) Distribution of T-cell colors by maternal and fetal origin, placental location, and CD4 and CD8 expression. (**E**) Distribution of CD4 T-cells colored according to the cell origin and overall expression of genes associated with Th1 and Th2 cells.

For detailed annotation of T-cells including resting and activated T-cells following annotation of previous attempts (*11*), selected 16,095 cells of T- and NK cells were subsequently reanalyzed, and eight types of T-cells were identified (Fig. 2C). From this point, the capitalized cell names \refer to the individual cell clusters we identified using scRNA-seq. For example, Tem_Trm_cyt_T indicates T effector memory tissue-resident memory cytotoxic T cells, and Tcm_Naive_hp_T indicates T central memory naïve helper T cell. We subsequently examined to those T-cells to determine whether the identified T-cell types are aligned with their known biological characters of T cells. As expected, Fig. 2 D and E support our confidence of the identified types T cells and the origin of the T cells. In Fig. 2D, the upper bound of the UMAP of T/NK cells mainly comprises fetal cells, and the type of cells identified are naïve cells, as presented in Fig. 2C. In the upper portion of these cell clusters, the expression of CD4 and CD8A genes exihibited mutually exclusive patterns and corresponded with those of CD4 helper T-cells and CD 8 cytotoxic T-cells, respectively. More specifically, CD4^+^ T-cells consisting of Tcm_Naive_hp_T (Tcm naïve helper T cell) and Tem_Eff_hp_T (T effector memory and Effector helper T cells) were scrutinized (Fig. 3E). Tcm_Naiv_hp_T clusters mainly consisting of fetal cells (upper part of Fig. 3E), whereas Tem_Eff_hp_T cells are maternal cells (others of Figure 3E). However, fetal, and maternal CD4^+^ T-cells are scattered throughout the placenta, suggesting broad interactions between semiallogenic cells in the placenta. Using the module scoring function of Scanpy, the overall expression of canonical genes of T helper (Th) cells, including Th1, Th2, Th22, Th17, Th9, and Tfh (T follicular helper T cells), were investigated to identify the subtype of T helper cells (*17*). The module score suggests that CD4^+^ T cells are Th1 and Th2 cells, which are known key players in pregnancy-associate immunity (*18*) (see table S2 for the list of genes associated with Th cells). Again, it allows us confidence in the data.

**Fig. 3.**
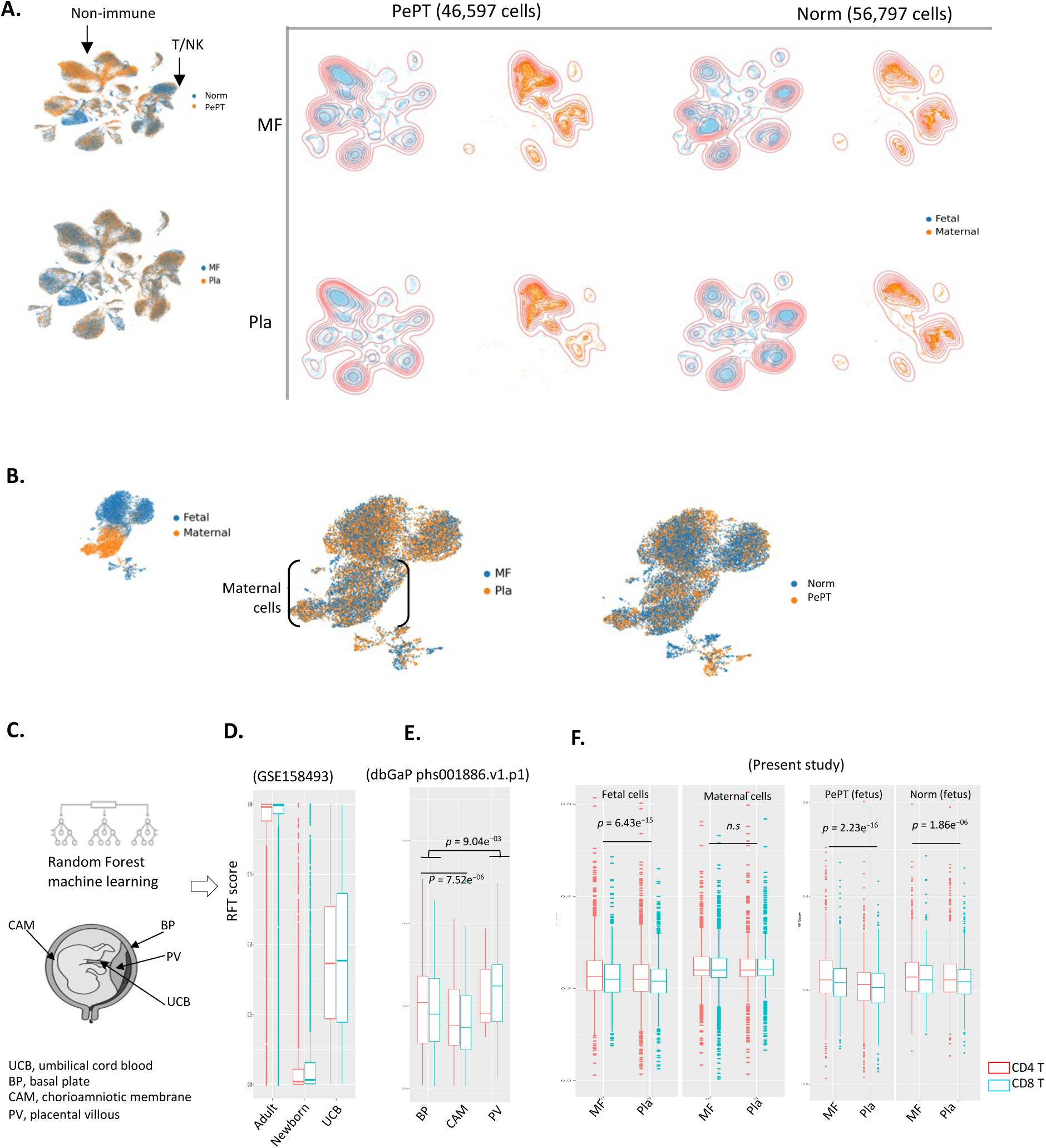
Interspersed distribution of semiallogeneic cells and immune transition of fetal T-cells. **(A)** Density projection of cells by placental layers and outcomes of pregnancy, where the colors of the dots indicate the origin of the cells. (**B**) Interspersed semiallogenic T/NK cells by placental unit and Norm and PePT pregnancy outcomes. (**C**) Schematic overview of the random forest (RF) model of our previous attempt (Bunis et al., 2020, *Cell Reports*) to grade the degree of T-cell maturity. (**D**–**E**). Validation of our RF model using T-cells from blood samples (GSE158493, *n* = 27) and cells from diverse placental regions from normal pregnant patients without labor (dbGAP phs001886.v1.p1, *n* = 9). (**F**) Results of RFTscores in our data from diverse placental layers (MR, Pla), origins of cells (fetal, maternal), and outcomes (Norm, PePT).

Taken together, our data well recapitulate the single-cell landscape of the human placental layers in favorable alignment with the known immunological characteristics, including representation of naive T-cells among fetal cells. Moreover, given the abundance of T/NK cells from semiallogenic origin within the placenta, our data seem promising for addressing the hypothesis of pregnancy-associated immune harmonization.

### Interspersed distribution of semiallogeneic cells throughout the placental layers

Based on our confidence in our data for a single-cell landscape of placental layers, we examined the suggested IIF model. In Fig. 2, a substantial fraction of maternal cells is observed in Pla, which undermines the central premise of the IIF model. We subequently scrutinize this further. The abundance of semiallogenic cells by placental region (i.e., Pla and MF) and pregnancy outcome (i.e., Norm and PePT) were compared using the density projection. Figure 3A presents the abundance of cells according to the corresponding origin (blue= fetal; orange=maternal cells), regions, and condition (i.e., Norm and PePT). Like T/NK cells, other nonimmune cells of maternal and fetal origins are spread throughout the Pla and MF region under both of Norm and PePT. Immune cells, including T and NK cells from different origins, are well intermingled across placental regions in both Norm and PePT conditions (Fig. 3B). In the fig. S1, our validation sample (PePT8) reproduces our observation that semiallogenic cells including nonimmune and immune cells are already intermingled. Therefore, in terms of frequency, the difference of degree of excessive interactions between semiallogeneic cells in Norms and PePTs is poorly supported by our observations.

Chronic inflammation is a notorious pathology of preeclampsia that impacts immunological balance (*19*). Although similar interactions between semiallogeneic cells throughout fetoplacental unit and both of PePT and Norm conditions are expected, it remains unclear whether proinflammatory triggers drive immune transition in PePT immune cells, disrupting immune tolerance during pregnancy. To examine the broad disruption of the immune adaptation of T-cells in PePT, we deciphered transition of T-cells by pregnancy outcomes and placental units. In a previous attempt, using the random forest (RF) model, we identified transition of T-cell (i.e., an adaptation of feto-maternal cell suppressing allograft rejection responses) that occurred gradually during the gestational period at the single-cell level (Fig. 3C) (*20*). The degree of T-cell transition measured in RF T score (RFTscore = [0,1], on y-axis). For example, a relatively larger RFTscore indicates more adult-like developmental staging of the immune system and a predisposition toward more adult-like inflammatory responses after stimulation, whereas lower scores suggest a more fetal-like origin and predisposition to tolerant. T-cells in umbilical cord blood showed an intermediate level of immune transition, whereas adult and fetal T-cell were converted into extreme tails of the RFTscore (GSE158493, N=27) (Fig. 3D). The RFTscore showed consistent results in another independent attempt involving of T cells from distinct placental regions including basal plate (BP, maternal region of placenta with higher RFTscores), placental villous (PV, middle layer), and chorioamniotic membrane (CAM, near fetus layer with low RFT) (p-value of t-test < 0.05) (*21*). Figure 3F shows differential RFT values in CD4^+^ and CD8^+^ T-cells from fetal origins across MF and Pla (p-value 6.43E-15). It is conserved in both PePTs (p-value 2.23E-16) and Norms (p-value 1.86E-06), which manifest that the overall disruption of immune transition in PePT is un-supported hypothesis. The RFT value in maternal T-cells across the MF and Pla were altered monotonically. The results in Fig. 3C-F suggest conserved immune transition of fetal T-cells, even under neighboring semiallogenic cells in both Pla and MF. Although the contribution of the difference in gestation between Norm and PePT is uncertain, even under the proinflammatory of PePT, the overall immune transition of T-cell aligns in order. Therefore, the failure of immune tolerance may be due to specific influencers rather than excessive interactions with semiallogenic cell.

We conducted differential gene expression analysis as described in the Materials and Methods (p-value of false discovery rate (FDR) adjust <0.05) to prioritize the transcriptional signatures of PePT, which strongly breaks immune harmonization. Fig. S2 and data S1 present the 7,987 differentially expressed genes (DEGs) from all cell cluster DEGs. Among those DEGs, PLCG2 is a known differential transcriptional signature of preeclampsia (*22*), identified as a top-ranked DEG repeatedly across diverse types of cells, including Tem_Eff_hp_T, Fibroblast, and Tcm_Naive_hp_T. Although other top-ranked DEGs across cell types recapitulated the results of previous attempts, including VEGEA association with preeclampsia(*23*), the DEGs are substantially overlapped across cell types. Immune cell type-associated gene signatures for PePTs are disregarded because of the design principle of the calculation of DEGs for comparing identical cell types.

### Machine learning-based identification of immune influencers of PePT

From this point, identifying transcriptional signatures among immune cells of PePT via DEG-based analytics is challenging. Therefore, we again pursed a divide-and-conquer approach, aming to develop a machine-learning approach that involves identifying the cell-type associated single-cell transcriptional signatures and tease out PePT-linked gene signatures. Using an XGBoost algorithm, a call type classification model established and subsequently used to classify Norm and PePT cells in a step-wise manner. Here, we postulated that the utilized gene signatures of each classification step, such as T-cells of PePT, are the immune influencers of PePT. Figure 4A illustrates the step-by-step classification acrhitecture of our model. By repurposing devCellpy(*24*), the classification model consisted in three layers. The first layer identifies cell types based on the trained signatures of genes. In the second layer, the model reclassifies those cells into either PePT or Norm in corresponding cells, such as a Normal Treg cell. In the third layer, the identified cell type information and outcome conditions are reclassified again into either of the placental layers (i.e., MF or Pla). Using Shapley additive explanations (SHAP) value evaluation(*25*), we ranked the contributions of transcriptional signatures in each layer, allowing us to prioritize immune influencers of PePT.

**Fig. 4.**
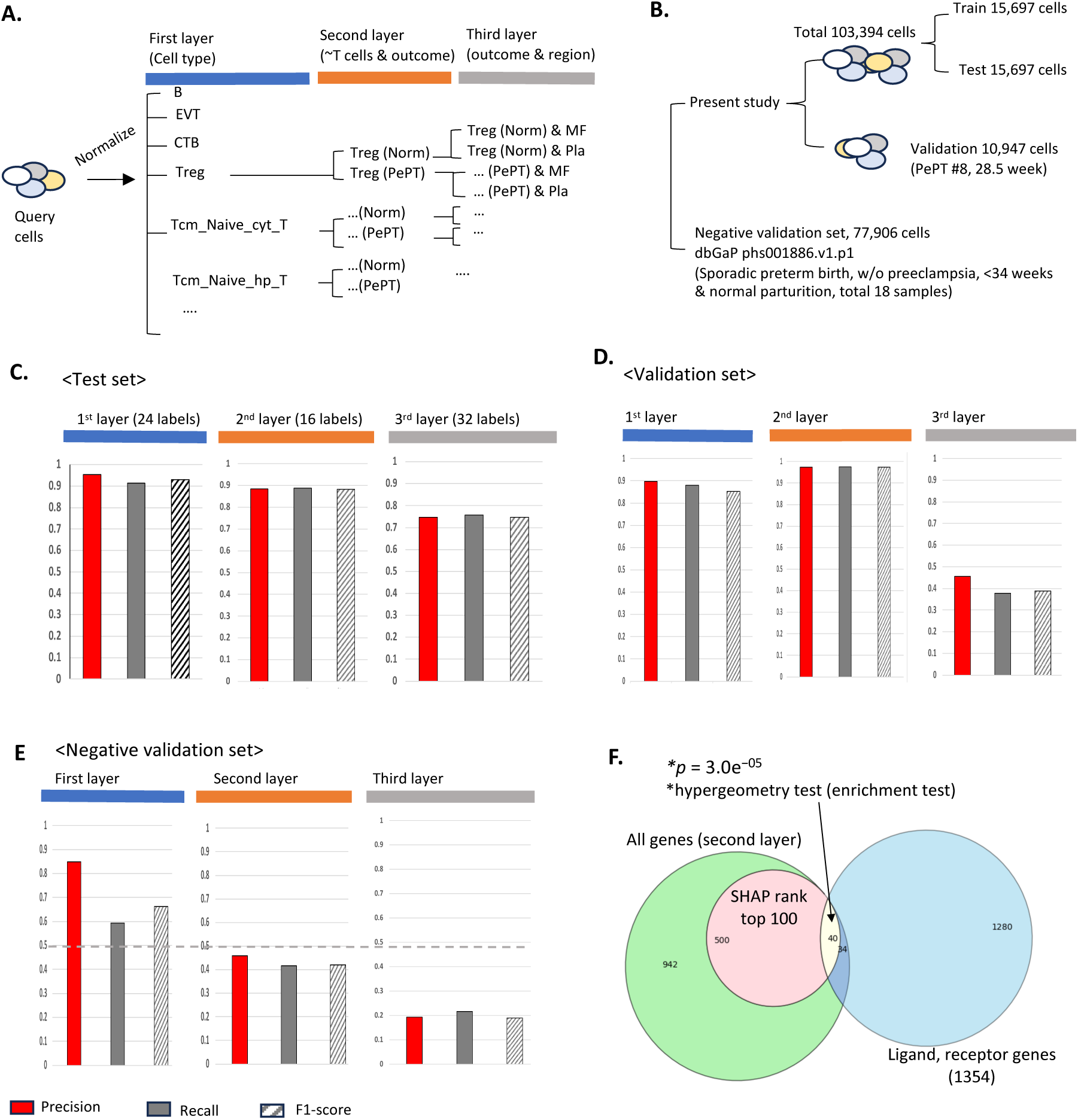
The machine learning approach involves interactions of immune cells as PePT- associated signatures. (**A**) Hierarchy of classification labels for devCellpy. Using an XGBoost approach, devCellpy allows the multilabeled and hierarchically structured identification of cells and assigned cell subtypes. In the first layer, the total number of classification labels is 24 cell types. In the second layer, devCellpy was subsequently established to identify immune cell types, including the Treg cells of PePT (Treg_PePT) and the Treg cells of Norm (Treg_Norm). In the third layer, our model classifies placental layers of cells in the second layer, such as Treg_PePT in MF (Treg_PePT_MF). (**B**) Performance evaluation methods, including twofold cross validation, use of independent validation set, and the negative set validation. (**C**) Average precision, recall, and F1-score across all the cell labels in the test set in the cross-validation settings. The total number of classification labels in each layer is presented. The detailed performance of each cell label in each layer is presented in figs. S3 and S4. (**D**) For the validation, we examined independent validation samples comprising the MF and Pla layers of a PePT patient (PePT #8, 28.5 weeks). The bar plot shows the average precision, recall, and F1-score for all the cell labels. (**E**) To examine the contribution of transcriptional signatures associated with gestation week differences, we evaluated our model using a negative validation set comprising cells from normal pregnancy and non-preeclampsia preterm parturitions. The bar plots show the average performance for each layer. The gray line indicates the prediction performance for random assignments. (**F**) Using the Shapley additive explanation (SHAP) rank value of the trained features of our model, we filtered the top 100 genes for each layer. Interestingly, these features are highly enriched with the ligand and receptor genes, indicating the contribution of immune cell interactions.

We evaluated the established model using an independent PePT validation sample that presented an untrained gestational week (28.5 week, PePT 8). The mean gestation of the training set is 31.6 weeks. As a more challenging problem, we employed a negative validation set to examine the trained devCellpy, which exhibited shortcut learning by focusing on gestational differences instead of identifying immune cell-associated transcriptional features. The negative validation set comprised 77,906 cells collected from placental tissues of normal pregnancies and under 34 weeks of gestations without preeclampsia (i.e., sporadic preterm birth; The database of Genotypes and Phenotypes (dbGaP, https://www.ncbi.nlm.nih.gov/gap) phs001886.v1.p1) (*21*) (Fig. 4B).

Figure 4C, 4D, and E4 show the performance of the developed XGBoost model in Fig. 4A. Because each classification layer covers over 16 up to 32 classification labels, figs. S3 and S4 present the detailed performance for each classification label, such as 24 of cell types in the first layer, and 16 types of PePT- and Norm-labeled immune cells in the second layer (Treg_Norm, and Treg_PePT). Within our test validation set, the established model shows successful performance across all layers (mean F1-score in the first layer = 0.93; second layer 2= 0.88) (details in fig. S3). In Fig. 4D, our model ensures consistent performance with the independent validation set in the first and second layers (i.e., mean F1-score in the first layer= 0.85; second layer= 0.97) (details in fig. S4). However, our model shows limited performance in the third layer in both the test set and validation sets (mean F1-score in the third layer in the test set= 0.74; in the validation set = 0.38). This finding highlights the disturbance of identifiying PePT-associated T-cell signatures in each placental region, as the results shown in Fig. 3F include differences in mature T-cell by their origin (i.e., adult maternal and fetal cells). Although there is underlying biological confounding in the third layer, the overall performance of our developed model allows us the confidence to identify immune cell influences of PePTs.

Moving forward, we addressed the possibility of shortcut learning of our model, as presented in Fig. 4B. The performance of our model in the first layer, the identification of cell types in non-preeclampsia preterm birth cells, was a favorable performance rather than random (mean F1-score in the first layer for all cell types = 0.66) (Fig. 4E; figure S4D and S4E). The underperformance of the established devCellpy model refutes the suspicion of shortcut learning, indicating that our model is promising for identifying immune influencers of PePTs (mean F1-score in the seconde layer 2 = 0.42; the third layer =0.19). We acknowledge that the utilized negative set included cells that underwent parturition labor, at least partially, and indicates the existence of T-cell activation-relevant perturbation for overall performance (*11*). However, the single-cell transcriptional signautres associated with labor are admixed with diverse cell types, including T-cells and other nonimmune cells including stromal cells (*26*). Taken together, confounding by the gestational period used for identifying differences in immune cells in the PePT condition is limited.

With confidence in our model, we investigated the main contributor of the model established to prioritize immune-associated influencers of PePTs. The SHAP analysis prioritized genes in the validated model. For example, the SHAP value ranks which one of the expressed genes mainly contributes to calssify Tregs of Norm and PePT. Figure S5 presents the top 20 of contributing genes for each layer of our classification model. Those top-ranked genes are aligned with known biological characteristics of each classification layer, such as CD3d for classifying T-cell groups (*27*) and COL1A1 for fibrogenesis of placental tissue (*28*). Interestingly, we revealed that the top 100 ranked genes, which contribute to identify immune cells of PePT, are highly enriched with ligand-receptor pairs using a comprehensive database (p-value of hypergeometry test = 3.0e-04.; Fig. 4F) (*29*). Among the 500 genes in the SHAP ranked in the top 100, fourty genes are overlapped with the database of ligand-recepter genes covering 1354 human genes. Thus, the types of cell-to-cell communication, including the T-cell types in the second layer (MAIT, Treg, Tcm_Naive_cyt_T, etc.), differ between PePTs and Norms. These 40 genes are prioritized as the immune influencers of PePTs. In data S1, we present a list of SHAP-ranked genes and prioritize 40 ligand-receptor pairs as immune influencers of PePTs.

### Validation of posed immune cell interactions in PePT

Though further analysis of the genes listed as immune signaturs of PePTs, we quantified the degree of communication between cells via those 40 ligands and recepters, including SPP1, PTGER, and CD44. Based on a comprehensive dataset of ligand recopters in human, compared with random gene expression, CellPhoneDB inferes the degree of interactions between cells based on coexpression of ligands and receptors (*29*). Figure 5A and 5B visualize identified ligand-receptor interactions in Norm (Fig.5A) and PePT (Fig, 5B) using a chord diagram consisting of arc blocks for cell types and chord lines for interactions between cells. We only drew chord lines between cell blocks with statistical significance (p-value of N-permutation < 0.05). The width and color of the chord lines depict the number of mean interactions between cell clusters. For instance, a thin, gray chord indicates relatively meager interactions between cell pair, whereas a thick, red chord means tight communication between cells.

**Fig. 5.**
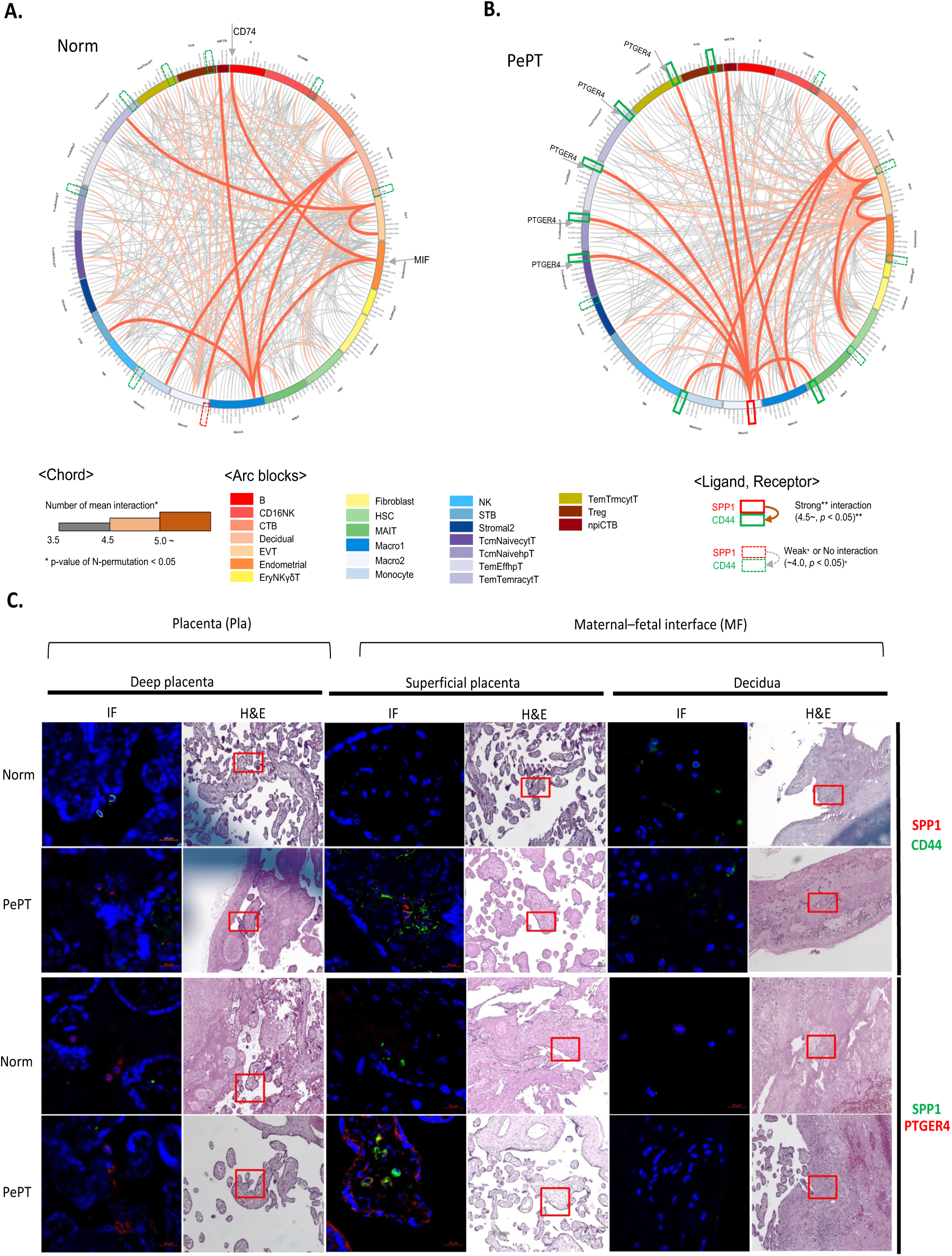
Differential interaction between SPP1 and CD44 in PePTs. **(A)** Chord diagram showing inferred interactions between cells from Norm samples. The color and width of the lines represent the strength of inferred interactions using CellPhoneDB (version 4). All visualized lines are statistically significant (*p* < 0.05, N-permutation test). Blocks of arcs present cell clusters. (**B**) Chord diagram showing inferred interactions between cells from PePTs. The green and red boxes indicate the SPP1 ligand and the CD44 receptor, respectively. (**C**) Experimental validation of identified differential interactions, including SPP1–CD44 and SPP1– PTGER4, using immunofluorescence (IF) and hematoxylin and eosin (H&E) staining. IF staining reveals the colocalization of SPP1 and CD44 proteins between cells. The H&E staining, particularly in the red boxed regions, highlights the association of inflammatory and immune response activity involving immune cells, such as T-cells.

In Norms, endometrial (like) and B-cells closely communicate via MIF (macrophage migration inhibitory factor) and CD74 (gray arrows in Fig. 5A). CD74 promotes the formation of an immunosuppressive tumor microenvironment by intracting with MIF (*30*). Thus, our results support a hypothesis that an immumesuppressitve mechnism is ahred between cancer and pregnancy (*31*). Interestingly, peculiar interactions between macrophage and diverse types of T-cells, including Treg, and Tcm_Naive_cyt_T were identified in PePT (Fig 5B). SPP1 (Secreted phosphoprotein 1), a gene known to be involved in cancer cell growth, tight interacts with T-cells via the CD44 receptor. Likewise, interactions via SPP1 and PTGER4 (a G-protein coupled receptor that can activate T-cell factor signaling) are also exclusively identified in PePTs. The green and red boxes highlight SPP1 (ligand) and CD44 (receptor), respectively. Figure S6 shows the identified cell-cell interactions in the corresponding placental regions, including the Pla and MF regions in the validation sample (PePT#8). Although the interactions between CD74 and MIF were relatively weaker than merged data of Figure 5A, those interaction still significant (p-value <0.05). The differential interactions of PePT consisting of SPP1 and CD44 (or PTGER4) are conserved whereas the validation sample (PePT#8) presented heterogenious interaction due to the limited cell numbers.

Figure 5A and 5B is consistent with our biological understanding, allowing us confidence inour analysis at in silico level. Using immunofluorecence (IF) and hematoxylin and eosin (H&E) staining, we examined the spatial colocalization of SPP1, CD44, and PTGER4 in each placental layer of Norm and PePT (Fig. 5C). IF staining demonstrated the localization of the SPP1, PTGER4, and CD44 proteins between cells. The placental layer comprises three pathological layers, including the deep placenta, superficial placenta and decidua. The superficial placenta and decidua region minaly corresponded mainly to the MF region, whereas the regions corresponded to Pla. In summary, SPP1 (red) and CD44 (green) protein-expressing cells are speicifcally colocalized in the MF of PePT, indicating tight communication. Moreover, the H&E staining, particularly in the red-boxed regions, highlights the associations with inflammation and immune response activity involving immune cells, such as T-cells. However, the colocalization of cells expressing SPP1 and PTGER4 in the MF of PePT remains open to interpretation in IF staining.

These results suggest that interactions between macrophage and T-cells via SPP1 (ligand) and CD44 (receptor) are specifically exist in placenta of preeclampsia.

### Proteomic differences between PePT and Norm aligned in the pathway of SPP1

Figure 5 manifests that PePT-specific local interactions between immune cells. The associations of SPP1 and CD44 with systemic inflammation in PePT during conception are unclear. Ought to identify systemic associations with PePT-specific cell-cell interactions, including SPP1 and CD44, we conducted Olink proteomic profiling using urine and plasma samples gathered via diverse gestational ages. As depicted in Fig. 6A, 30 non-invasive samples (i.e. urine, and plasma) were collected from total 30 women consisting. Because CD44 is a nonsecreted protein, we aligned the corresponding genes to comprehensive pathways in KEGG (*32*) to identify perturbed downstream secreted proteins. Although SPP1 is a secreted protein, 189 markers in the Olink panel showed non-intersection. Those 189 proteins constitute the union set of three Olink panels which consists of 96 Immune response, 97 Inflammation, and 48 cytokine panels. Thus, among the markers detected through Olink proteomics, we compared the abundance of downstream proteins associated with SPP1 and CD44 in the urine and plasma samples of the Norm and PePT groups.

**Fig. 6.**
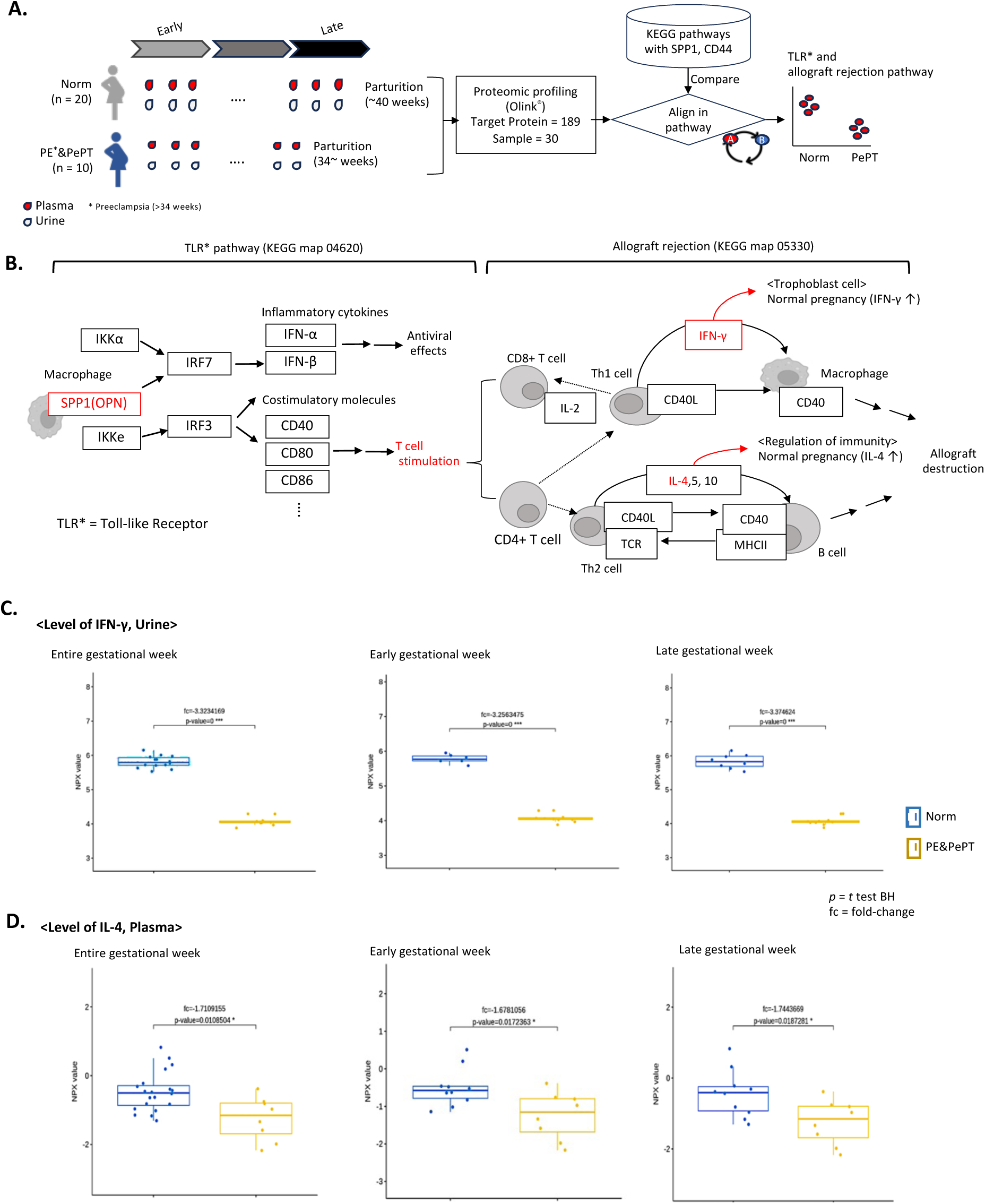
Differential expressed proteins (DEPs) in PePTs. (A) Analysis of proteomic signatures from noninvasive samples in Norm and preeclampsia and PePTs. (**B**) Prioritized pathways of KEGG that are aligned with SPP1 or CD44. (**C**–**D**) Proteomic expression levels of IFN-γ and IL-4 in the urine and plasma from Norm and PE&PePT patients by trimester. All urine and plasma samples were prepared as longitudinally matched sets across gestation from each patient. Normalized protein expression (NPX) values in the *y*-axis are the units of relative protein concentration used in the Olink proteomics analysis.

Figure 6B presents an overview of prioritized pathways that harbor SPP1 and CD44. Out of 576 pathways of KEGG (release 112.0, October 1, 2024), the Toll-like receptor (TLR) pathway (map04620) was selected by overlapping with SPP1, and then the pathway of allograft rejection (map05330) was prioritized as a downstream path by aligning the T-cell stimulation module on the TLR pathway. Notably, although SPP1 (also known as osteopontin, OPN) is a multifunctional protein, it is widely involved in tumor progression, such as by inhibiting T-cell activity and promoting metastasis through interactions with CD44 and T-cells (*33*). As a downstream pathway of T-cell stimulation of TLRs, allograft rejection pathway involves several target proteins in our Olink profile, including inteferon-gamma (IFNγ) and interleukin-4 (IL-4). In normal pregnancy, IFNγ plays a critical role in the proinflammatory cascade triggering angiogenesis of placental tissue (*34*). The anti-inflammatory cytokine, IL-4, is also essential for successful pregnancy because it activates Treg and Th2s, inducing immune tolerance of the fetus(*35*).

Considering the biological roles of these target proteins in the Olink profile, we examined the expression levels of secreted proteins from urine and plasma of pregnant women before parturition. Overall, the typical multidimensional scaling (MDS) dimensional reduction techinque (*36*) for Olink profiling of immune responses, proteins from patients are closely clustered regardless of patient group or time of gestation. However, the cytokine and inflammation panels present distantly clustered view, incidating differences in cytokines and inflammation proteints by early and late gestation (fig. S7A, S7B and S7C). Although the proteins profiled in our Olink panel are partial (total of 189 secreted proteins), 96 inflammation and 48 cytokine panels are appropriate to revealing proteomic differences by getational age and pregnancy outcomes including PE&PePT. Therefore, we have focused more on downstream pathways including cyokine secresions and Inflammation associated proteins in the Olink.

Downstream proteins of SPP1 (OPN) including IFNγ and IL-4 presented significantly decreased expression levels in patients with PE& PePT patients (p-value of t-test with Benjamini-Hochberg adjustment < 0.05; Fig. 6C and 6D). Both proteins are involved in 96 inflammation and 48 cytokine panels (data S1). The blue boxes and dots represent the levels of normalized protein expressions (Normalized Protein eXpression, NPX value) in Norm, whereas the dark-yellow dots are the NPX values of PE&PePTs. The differential expressions of IFNγ and IL-4 between Norm and PE&PePT is constant in both of the early and late trimesters, indicating that the associated interactions between SPP1 and CD44 are initiated in early pregnancy. Although evidence for causal relationships between SPP1-triggered immune responses and identified DEPs of cytokines, including IFNγ and IL-4 is lacking, MDS of 96 immune response panels already manifests the homogeneous secretion proteomic profiles between the Norm and PE&PePT. Thus, differential proteomic levels of IFNγ and IL-4 in the plasma and urine of patients with PE&PePT are associated with SPP1 expression in macrophages in the placenta.

In conclusion, in a preeclampsia placenta, the local interactions between macrophages and T-cells via SPP1 ligands and CD44 receptors influence downstream cascades, including the allograft rejection pathway, which disrupts systematic immune tolerance associated with differential proteomic levels of IFNγ and IL-4 in the plasma and urine. Owing to the detection of these differences across gestation, those noninvasive markers appear promising for clinical applications.

## Discussion

To unravel the immunological phenomenon of pregnancy,we investigated large-scale single-cell transcriptomic profiles in Norm and PePT patients while considering placental regions, including the Pla and MF. Although PePTs are induced during clinical intervention at approximately 34 weeks of gestation, we defined PePTs as severe cases of preeclampsia (PE). We began under the hypothetical frame for the failure of normal fetomaternal tolerance of PePT by considering either the excessive semiallogeneic interaction of cells (i.e.m IIF model), or preeclampsia-specific cellular interaction (i.e., IT model). The premise of the IIF model is that the physical segregation of semiallogenic cells during placental development leads to immune toloerance. However, the wide dispersion of semiallogenic cells in placenta, including immune-associated T-cells for both Norm and PePT undermines IIF model. Moreover, mature the degree of maturity of fetal T-cells of fetus gradually shifts the trajectory toward an adult-like state across placental layers, whereas fetal cells adjacent to maternal cells, challenging the premise of the IIF model again. The premise of the IT model is the existance of differential interactions of cells between Norm and PePT. Using the XGBoost approach (*24*), we successfully developed a classifiction model for Norm and PePT immune cells which presented prioritized differential transcriptional signatures of PePTs at the single-cell level. Notably, 500 gene prioritized expressions for PePT immune cells were highly enriched with human ligand and receptor genes, indicating the existence of influencers of PePT under the IT model (*p-value* of hypergeometry test = 3.0e-05). We filtered the interactions of those genes uing CellPhoneDB (*29*), then examined using IF and H&E staining. The specific immune cell interactions between SPP1 and CD44 are presented exclusively in the PePT placenta MF layer. Protein downstream of SPP1 (OPN), including IFNγ and IL-4, showed significantly decreased expression levels in patients with PE&PePT. Local interactions between macrophages and T-cells via SPP1 would influences the systemic immune tolerance of PePT. Exploring single cells via our divide-and-conquer approach clearly unravelled that PE is not complete failure of immune balance but instead undergoes fractional breakdown of immune coordination throughout gestation.

Here, we showed that, as in the tumor, an immunosuppressive microenvironment is formed in the normal placenta. For example, MIF binding to CD74 induces a signaling cascade resulting in the regulation of naïve B-cells for tumor development (Fig. 5) (*30*). Norm and PePT placental cells commonly interact with the decidua and macrophages via SCAR5 and FTL, which are associated with tumor metastasis and growth (*37*). Although the presented single-cell landscape showed a distinct composition of cells with a tumor microenvironment, results of Fig. 5A and 5B shows mechanisms of immune tolerance shared between pregnancy and cancer. Owing to the invasion phenotype of trophoblast cells, previous studies have reported shared immune mechanisms between cancer and early trimester samples (*38*). Based on the final trimester cells, our analysis of cell-cell interactions suggested that the immune mechanisms shared between pregnancy and cancer continue throughout pregnancy. Our analysis recapitulates that the immune suppressive mechanism found in cancer continues from the initial establishment of trophoblast cells to the third trimester. The knowledge gained from analyzing the similarities between the state of pregnancy in terms of single-cell gene expression and the pathological state of cancer could lead to the identification of new potential targets for PE.

The present study has several limitaions. Single-cell level gene expression analysis alone may not completely capture systematic inflammation; however, establishing a transcriptomic foundation for PePT and identifying the specific cell types and those interactions involved hold substantial value for future studies. Furthermore, machine learning and bioinformatic techniques can provide robust inferences regarding PePT-specific genes and the participating ligands and receptors of immune cells. Regardless, we acknowledge that additional validation of the association between SPP1 and identified proteomic differences, including those of IFNγ and IL-4, is required to verify their causality. In addition, owing to the limited resolution of current spatial biology technology(*39*), interactions between fetomaternal cells are remained unclear. Therefore, any potential contribution of interactions between semiallogenic cells and the inflammatory case conditions described here cannnot be excluded. Finally, our findings also suggest that our predicted single-cell model of the placenta, and proteomic signatures from the plasma and urine of pregnant women should be tailored to the target population, given our observation that ethnicity may be a contributing factor. This caveat further underscores the importance of custamizing biomarkers of obstetrical diseases.

In conclusion, the findings generated here provide insight into immune signaling via a specific pair of influencers, SPP1 (ligand) and CD 44 (receptor), during pregnancy underwenting PE. Our findings highlight the utility of machine learning and single-cell technologies for shedding light on cell type-associated interactions that are specifically implemented in the PePT placenta and indicate the potential use of placenta-derived single-cell signatures for developing noninvasive biomarkers to predict PePT, a leading cause of neonatal mortality worldwide.

## Materials and Methods

### Study design

This prospective cross-sectional study included women who delivered at normal term (Norm) (≥37 weeks of gestation, n = 3), who had preeclampsia with severe features, and who had preterm preeclampsia (PePT) (::36 weeks of gestation, n = 4). All women underwent elective cesarean delivery. To avoid a marginal overlap of gestational weeks between Norm and PePT, we applied a strict threshold for selecting PePT cases (::34 weeks of gestation). Other patient samples, including plasma urine, were collected from the biobank of Konyang University Hospital. The Institutional Review Board of Konyang University Hospital reviewed and approved the study design.

### Analysis of droplet-based single-cell RNA sequencing data

Placenta tissues were obtained during cesarean section. The tissues of the MF and PL layers of the placenta were freshly isolated, and the samples subsequently underwent general cell processing steps, including library preparation and sequencing. In brief, cell viability and concentration were measured using a Countess II FL Automated Cell Counter (Thermo Fisher Scientific, USA). An RNA-seq library was prepared using a Chromium Single Cell 3′ Library (10x Genomics) according to the manufacturer’s instructions. Samples (>10,000 cells per sample per condition) were loaded and sequenced on an Illumina HiSeq-x sequencer, with an average of 11117.7±3057.9 cells per sample. The sequencing data were aligned and quantified using the Cell Ranger Single-Cell Software Suite (10xGenomics) against the *Homo sapience* genome (GRCh38) provided by Cell Ranger. Using Scanpy (version 1.9.1) package of Python (*40*), cells with fewer than 200 detected genes, fewer than 1000 UMI counts, cells with the total mitochondrial gene expression exceeded 20%, and cells with more than 7000 genes expressed per cell were removed. All of raw read data were adjusted using SoupX to minimize noise of ambient RNA(*41*). Doublet cells are filtered based on the results of Scrubelt (*42*). To normalize the data, global scaling, analysis of highly variable genes, and dimensionality were reduced using principal component analysis (PCA), which was conducted before UMAP was obtained to visualize cell clusters effectively (*43*). Following the initial quality control and preprocessing steps, we merged all the scRNA-seq data of each tissue and individual. Harmony (*44*) was used as a batch effect correction algorithm after the clustering results of BBKNN were compared (*45*). Similarly, in CellTypist (*46*), using the scleto2 package we developed (https://pypi.org/project/sceleto2/), we identified cell types using a logistic regression modeling approach based on the reference label of cells in the human placenta in late trimester (*21*). Owing to the absence of low level T-cell annotation in those late trimester data, the reference model of Celltypist (Immune_All_Low) was utilized for the detailed labeling of T-cells, including Treg.

### Identification of fetal and maternal cell

We applied a dual-cross confirmation approach to identify the origin of the obtained cells, including fetal and maternal cells in a given placenta. First, we utilized a method to cluster cells by genotype using Souporcell(*15*) and then classified the cells into two genetically distinct clusters. Second, to verify the origin of each cluster, genotypic profiles derived from Souporcell were matched against the genotype variance data of maternal peripheral blood mononuclear cells (PBMCs) and fetal cord blood, acquired using an Illumina GSA v3 genotype array. Single nucleotide polymorphisms from the clusters were aligned with those from the maternal PBMC genome to evaluate concordance levels. The cluster showing the greatest alignment with the maternal PBMC genotype was labeled the maternal cell cluster, and the other cluster was identified as fetal. In brief, we examined over 1000 alleles (mean 1115.875± 381.2 per sample) to confirm cells from genetically distinct clusters.

### Machine-learning model and analysis of ligand-receptor interactions between cells

To identify critical features associated with Norm and PePT conditions, we applied devCellPy, a hierarchical machine learning framework based on XGBoost (*24*). This classification model was structured into three levels: the first level categorized broad immune cell types, the second level focused on T-cell subtypes differentiated by Norm and PePT conditions, and the third level identified regional variations within MF and PL layers. We prioritize contribution of genes of the validated model using SHAP (Shapley Additive Explanations) analysis, then determined that top-ranked genes highly enriched with ligand-receptor genes in the comprehensive human ligand, receptor databased of CellPhoneDB(*29*). The statistical significance of enrichment of the ligan and receptors are determined using p-value of hypergeometry test (*p* < 0.05). To identify differences in cell–cell communication between the Norm and PePT conditions, we used CellPhoneDB (version 3).

### Immunofluorescence (IF) staining

Antigens were retrieved from fixed tissue sections using citrate buffer (pH 6.0), followed by heating to 60°C overnight. Nonspecific binding sites on the sections were blocked with 5% normal goat serum in PBS for 1 hour at room temperature to prevent nonspecific antibody binding. Primary antibodies diluted in an antibody diluent solution were applied overnight at 4°C. After washing, fluorescently labeled secondary antibodies were applied for 1 hour at room temperature. The sections were washed again and mounted with a medium containing 4′,6-diamidino-2-phenylindole (DAPI) for nuclear staining. Negative controls were prepared by omitting the primary antibody, while positive controls were included to verify staining specificity. The fluorescent signals were detected using a Carl Zeiss LSM700 confocal microscope, and the staining intensity was analyzed using Zeiss Zen software.

### Tissue collection and preparation for IF staining

Placenta tissues were obtained from patients’ post-partum. These tissues were fixed in 10% neutral buffered formalin, then embedded in paraffin. Sections were sliced to a thickness of 5 µm using a microtome and subsequently stained with Hematoxylin and Eosin (H&E) for histological analysis.

## Data and code availability

The scRNA-seq data were deposited at GEO and are publicly available as of the date of publication. The accession numbers are listed in the key resources table. This paper does not report the original algorithm code. Other source codes for the figure vignette and genotype analysis from the results of Souporcell are presented (https://github.com/hypaik/PePT_vignette). Any additional information required to reanalyze the data reported in this paper is available from the lead contact upon request.

### Olink proteomic profiling

Patient samples, including plasma and urine with PE, PePT and Norm controls, were enrolled at the Biobank of Konyang University Hospital. We selected 80 urine and plasma samples from 40 women in their early and late trimesters in a pairwise manner. All samples analyzed were collected before parturition. We selected 80 urine and plasma samples from 40 women in their early and late trimesters in a pairwise manner. All samples analyzed were collected before parturition. To randomize the samples, we prepared experimental source plates in an order appropriate for the run plate layout, randomized the samples, and arranged them appropriately. The Olink Target panel process was executed according to the manufacturer’s protocol. The antibody pairs with attached DNA tags were added to the serum/plasma samples(1 μL) and allowed to bind to their target proteins overnight at 4°C for 20 hours. On the following day, the extension and amplification steps take place. Oligonucleotides brought into proximity were hybridized and extended using DNA polymerase. This newly created piece of DNA barcode was amplified by polymerase chain reaction (PCR). The thermocycler conditions were 50°C for 20 min and 95°C for 5 min, followed by 17 cycles of 95°C for 30 s, 54°C for 1 min, and 60°C for 1 min. The final PCR step was maintained at 10°C. The final detection step quantifies the DNA reporters for each biomarker using high-throughput real-time quantitative PCR on the Olink Signature Q100 system. Raw data from Olink Signature Q100 were imported, validated to assess its quality, normalized, and analyzed statistically with Olink NPX Signature software. The cycle time (Ct) for each protein in each sample was obtained using an Olink Signature q100 instrument following the Olink panel experiment. These Ct values were normalized using interplate control (IPC) normalization to obtain normalized protein expression (NPX) via the Olink NPX signature software. In the quality check steps, samples were flagged as QC warnings based on deviation from the internal control median NPX value. Protein assays were filtered by applying a missing frequency, which depended on the limit of detection (LOD). Data processing and statistical analyses were performed using the OlinkAnalyze R package (https://cran.r-project.org). Differentially expressed proteins (DEPs) were determined using a raw p < 0.05 from a paired t test in which the null hypothesis was that no difference exists among groups. FDR was controlled by adjusting p using the Benjamini–Hochberg algorithm. All the data analysis and visualization of the DEPs were conducted using R software (version 4.2.2, www.r-project.org).

## Acknowledgement

This work was supported by the Korea Institute of Science and Technology Information (KISTI) (K-23-L02-C04-S01). This work was also supported by the Health Industry Development Institute (KHIDI HI22C1318, N-23-NT-CA01-S01). This research was also supported by the project of Building and Operating the Korea-Bio Data Station (K-BDS) Platform building and tool development (NRF-2021M3H9A2030520, N-23-NM-CA01-S01). This work was supported by the National Supercomputing Center with supercomputing resources including technical support KSC-2023-CRE-0526. M. Sirota was supported by UCSF March of Dimes Prematurity Research Center. This research was also supported by the Basic Science Research Program through the National Research Foundation of Korea (NRF), which is funded by the Ministry of Education (NRF-2017R1A6A1A03015713) and Konyang University Myunggok Research Fund (2020–3).

## TABLE OF CONTENTS FOR SUPPLEMENTARY MATERIALS

**Table S1.** Single cell RNA-seq data statistics

**Table S2.** Set of genes for the module scoring of Th cells

**Fig. S1**. Single-cell-level overview and density by placental layers in our validation sample (PePT8).

**Fig. S2.** Volcano plots of differential expressed genes (DEGs) of PePT and Norm

**Fig. S3.** Detailed performance of our established classification model in our test dataset.

**Fig. S4.** Detailed performance of established our classification model in the validation and negative sets.

**Fig. S5.** SHAP ranked genes in our classification model in layer 1 and 2.

**Fig. S6.** Differential interaction between SPP1 and CD44 by placental regions and validation set.

**Fig. S7.** Multidimensional Scaling (MDS) by placental regions and validation set.

**Data S1.** Total list of DEGs

**Data S2.** Target proteins in Olink panels.

## Funding

Korea Institute of Science and Technology Information (KISTI) (K-23-L02-C04-S01), the Health Industry Development Institute (KHIDI HI22C1318, N-23-NT-CA01-S01), the project of Building and Operating the Korea-Bio Data Station (K-BDS) Platform building and tool development (NRF-2021M3H9A2030520, N-23-NM-CA01-S01), the National Supercomputing Center with supercomputing resources including technical support KSC-2023-CRE-0526, UCSF March of Dimes Prematurity Research Center, the Basic Science Research Program through the National Research Foundation of Korea (NRF, NRF-2017R1A6A1A03015713), and Konyang University Myunggok Research Fund (2020–3).

### Author contributions

Hyojung Paik: Conceive the main idea, data acquisition, draft write, bioinformatic analysis design, supervise overall project, prepare all figures, and funding acquisition.

Tae Lyun Ko: Carried out bioinformatic analysis, data acquisition, supporting figures. Myungsun Park: Supporting bioinformatic analysis for data preprocessing

Jong-Eun Park: Supporting bioinformatic analysis

Danial Bunis: Supporting bioinformatic analysis for classification modeling

Marina Sirota: Supporting bioinformatic analysis for classification modeling

Byung Soo Lee: Carried out experimental validation

Hyeong-Sam Heo: Carried out experimental validation

Sung Ki Lee: Carried out sample preparation, supervise overall project, and funding acquition.

### Data and materials availability

GEO 290578

### Funding information

The funder has played no role in the research design, analysis overall, and writing.

