## SupplementalData for "Searching for influencers among placental immune cells in preeclampsia": table S1.docx

**Table S1. Single cell RNA-seq data statistics**

| **No.** | **Cluster cell label** | **Total cells** | **Fetus cells** | **Maternal cells** | **MF** | **Pla** | **Norm** | **PePT** |
| --- | --- | --- | --- | --- | --- | --- | --- | --- |
| 0 | Macro2 | 12638 | 12549 | 89 | 6346 | 6292 | 4286 | 8352 |
| 1 | NK | 9231 | 8363 | 868 | 3845 | 5386 | 7306 | 1925 |
| 2 | Stromal2 | 9058 | 9027 | 31 | 4130 | 4928 | 5405 | 3653 |
| 3 | Macro1 | 7723 | 135 | 7588 | 3897 | 3826 | 3650 | 4073 |
| 4 | Tcm_Naive_hp_T | 6596 | 6332 | 264 | 2486 | 4110 | 4721 | 1875 |
| 5 | EVT | 5601 | 5573 | 28 | 4146 | 1455 | 2427 | 3174 |
| 6 | Endometrial(like) | 5509 | 5501 | 8 | 5191 | 318 | 5195 | 314 |
| 7 | Monocyte | 5463 | 1594 | 3869 | 2932 | 2531 | 2220 | 3243 |
| 8 | Macro1 | 5126 | 1985 | 3141 | 2737 | 2389 | 1165 | 3961 |
| 9-0 | Tem_Trm_cyt_T | 1637 | 71 | 1566 | 884 | 753 | 1206 | 431 |
| 9-1 | Tem_Eff_hp_T | 1123 | 268 | 855 | 560 | 563 | 778 | 345 |
| 9-2 | Tem_Temra_cyt_T | 708 | 8 | 700 | 417 | 291 | 503 | 205 |
| 9-3 | CD16_NK | 610 | 8 | 602 | 326 | 284 | 353 | 257 |
| 9-4 | Treg | 474 | 56 | 418 | 265 | 209 | 320 | 154 |
| 9-5 | MAIT | 405 | 10 | 395 | 220 | 185 | 321 | 84 |
| 9-6 | Tcm_Naive_hp_T | 67 | 35 | 32 | 35 | 32 | 46 | 21 |
| 9-7 | Tcm_Naive_cyt_T | 9 | 9 | 0 | 3 | 6 | 5 | 4 |
| 10 | Macro2 | 3837 | 3810 | 27 | 2010 | 1827 | 1390 | 2447 |
| 11 | Tcm_Naive_cyt_T | 3452 | 3153 | 299 | 1211 | 2241 | 2674 | 778 |
| 12 | npiCTB | 2975 | 2967 | 8 | 1725 | 1250 | 1506 | 1469 |
| 13 | Monocyte | 2815 | 1657 | 1158 | 1150 | 1665 | 1514 | 1301 |
| 14 | NK | 2366 | 1478 | 888 | 1247 | 1119 | 727 | 1639 |
| 15 | EVT | 1882 | 1865 | 17 | 1189 | 693 | 748 | 1134 |
| 16 | NK | 1760 | 227 | 1533 | 927 | 833 | 1365 | 395 |
| 17 | B | 1629 | 1211 | 418 | 845 | 784 | 960 | 669 |
| 18 | HSC | 1560 | 1309 | 251 | 784 | 776 | 953 | 607 |
| 19 | Decidual | 1229 | 33 | 1196 | 939 | 290 | 765 | 464 |
| 20 | Fibroblast | 1173 | 1020 | 153 | 611 | 562 | 307 | 866 |
| 21 | CTB | 1162 | 1160 | 2 | 680 | 482 | 783 | 379 |
| 22 | Ery_NK_γδT | 1014 | 862 | 152 | 454 | 560 | 630 | 384 |
| 23 | Monocyte | 958 | 932 | 26 | 343 | 615 | 579 | 379 |
| 24 | STB | 892 | 866 | 26 | 378 | 514 | 474 | 418 |
| 25 | HSC | 664 | 660 | 4 | 352 | 312 | 317 | 347 |
| 26 | NK | 632 | 375 | 257 | 347 | 285 | 303 | 329 |
| 27 | B | 567 | 494 | 73 | 323 | 244 | 350 | 217 |
| 28 | Macro1 | 419 | 43 | 376 | 243 | 176 | 177 | 242 |
| 29 | EVT | 174 | 174 | 0 | 174 | 0 | 174 | 0 |
| 30 | NK | 116 | 78 | 38 | 51 | 65 | 88 | 28 |
| 31 | Stromal2 | 64 | 64 | 0 | 2 | 62 | 63 | 1 |
| 32 | Macro2 | 28 | 26 | 2 | 18 | 10 | 5 | 23 |
| 33 | Endometrial(like) | 24 | 1 | 23 | 16 | 8 | 21 | 3 |
| 34 | HSC | 24 | 23 | 1 | 6 | 18 | 17 | 7 |
|  |  | 103394 | 76012 | 27382 | 54445 | 48949 | 56797 | 46597 |
