## SupplementalData for "Searching for influencers among placental immune cells in preeclampsia": table S2.docx

**Table S2. Set of genes for the module scoring of Th cells**

| **Th cell name** | **Gene name** | **Official gene symbol used** |
| --- | --- | --- |
| Th1 | T-bet, IFN-γ, TNF-α, IL-2 | TBX21, T-bet, IFNG, IFG, TNF, IL-2, IL2, TCGF |
| Th2 | GATA3, STAT5, IL-4, IL-5, IL-13 | GATA3, STAT5, STAT5A(B), IL4, IL5, IL13 |
| Th17 | ROR-γ, IL-17, IL-22 | IL17, IL17F, IL22 |
| Th9 | PU.1, IL-9 | PU1, SPI1, IL9, HP40 |
| Th22 | AHR, BNC2, FOXO4, IL-22 | AHR, BNC2, FOXO4, IL-22 (IL22) |
| Tfh (T follicular helper) | Bcl-6, IL-21 | Bcl-6 (BCL-6, BCL6), IL-21 |
| iTreg (induced regulatory T) | IL-10, FoxP3, TGF-β | IL10, FoxP3, TGF-b(B), TGFB1 |
