## Supplementary figures and images for "Searching for influencers among placental immune cells in preeclampsia"

### FigS1_300.jpg

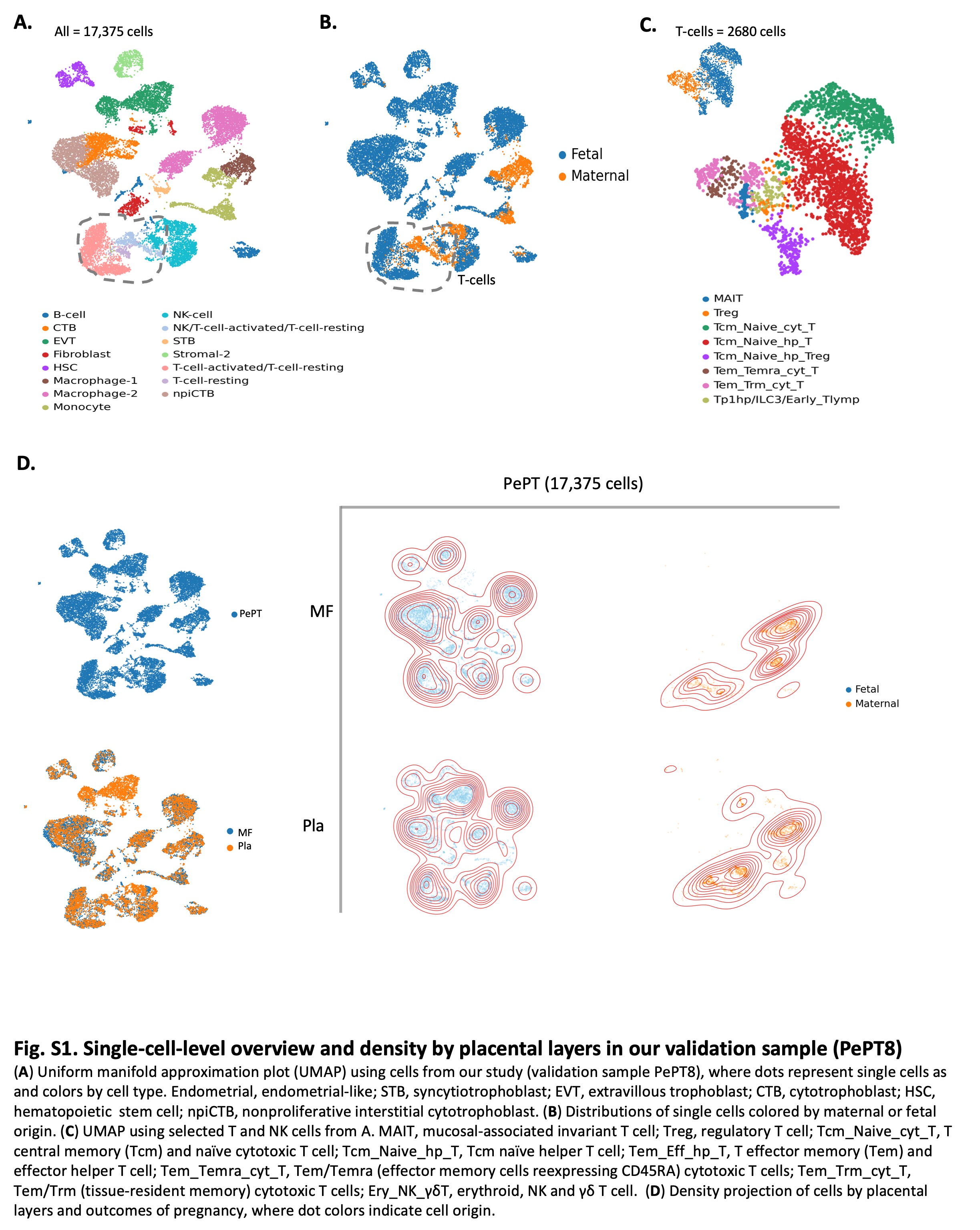

### FigS2_300.jpg

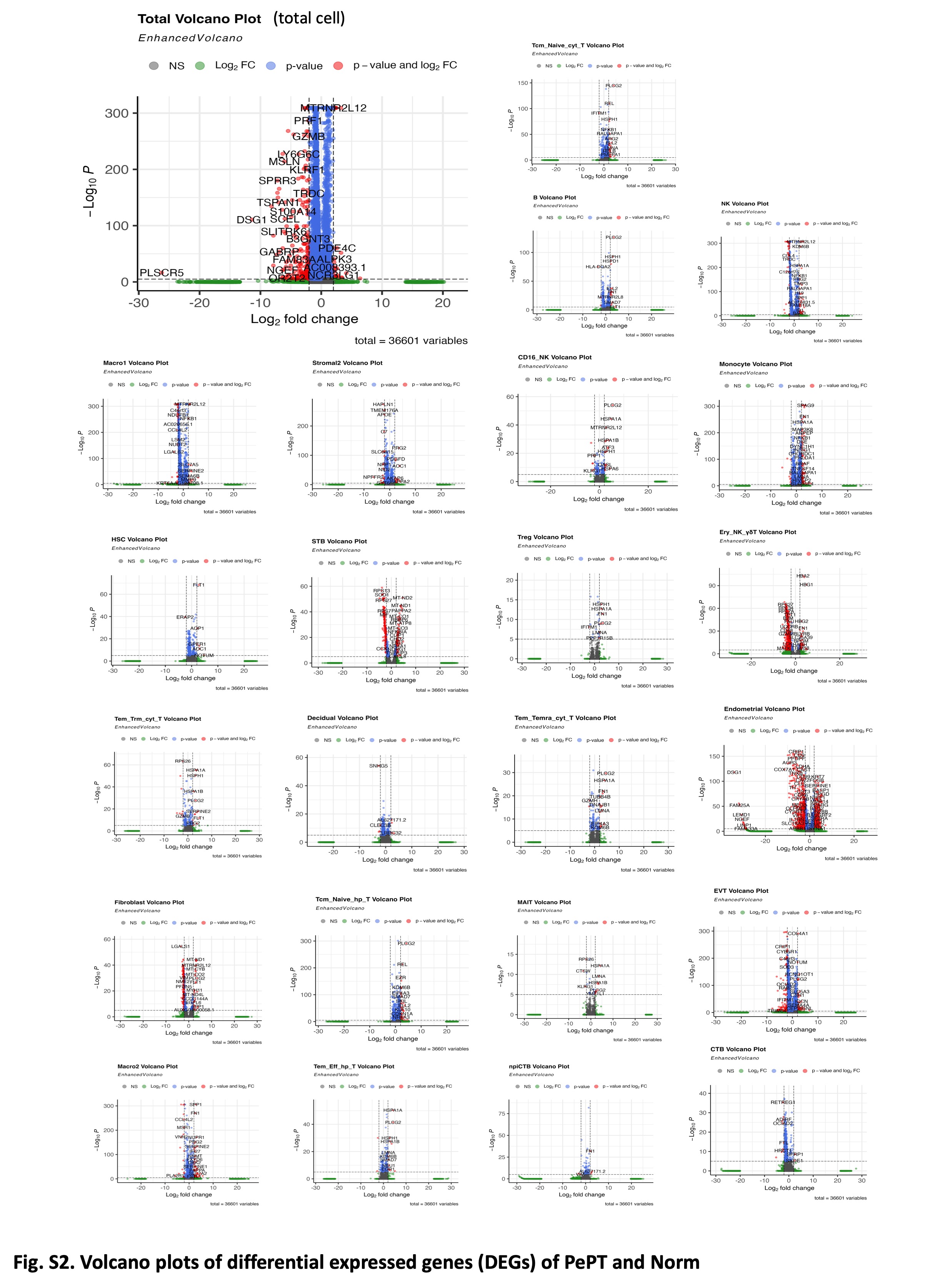

### FigS3_300.jpg

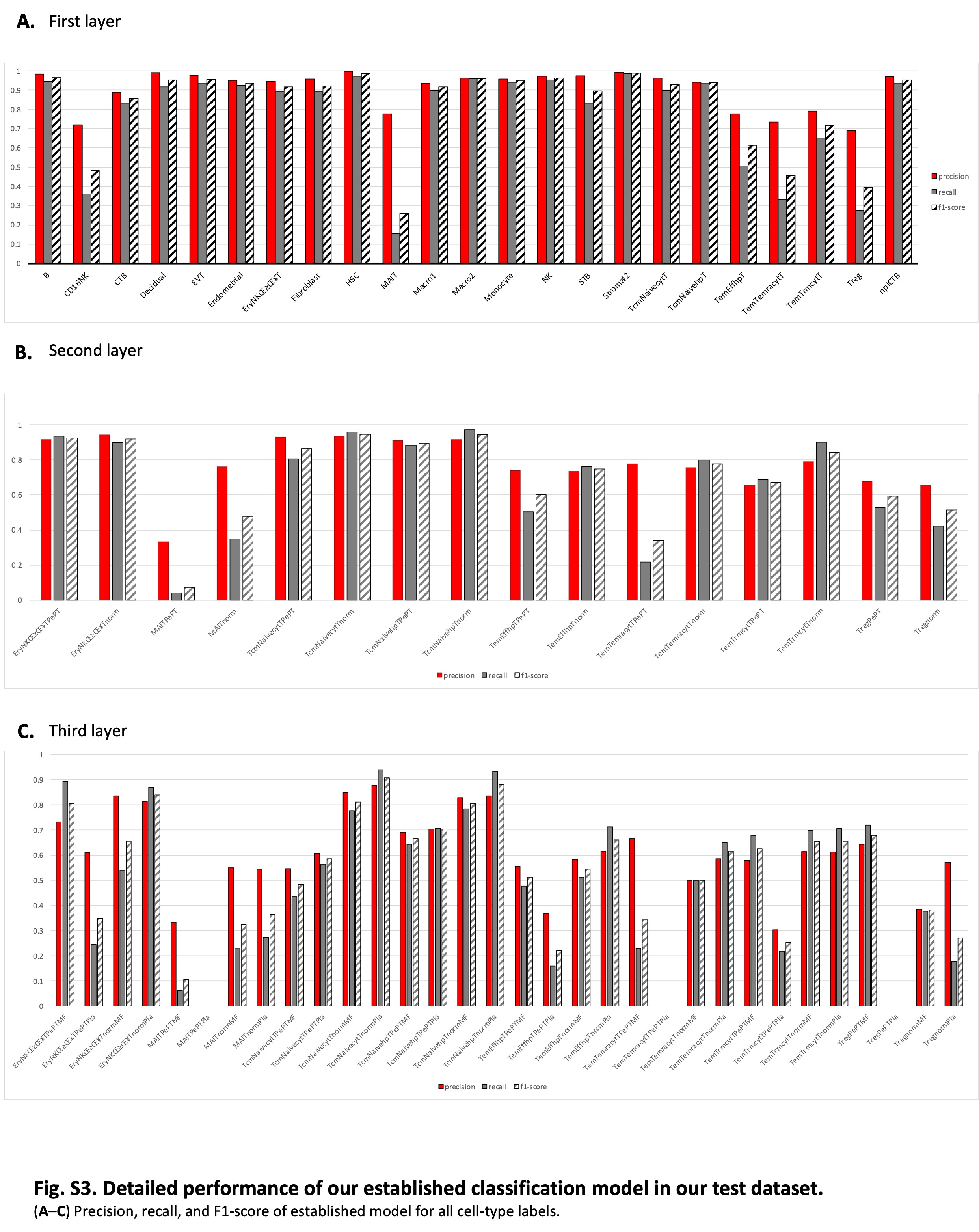

### FigS4_300.jpg

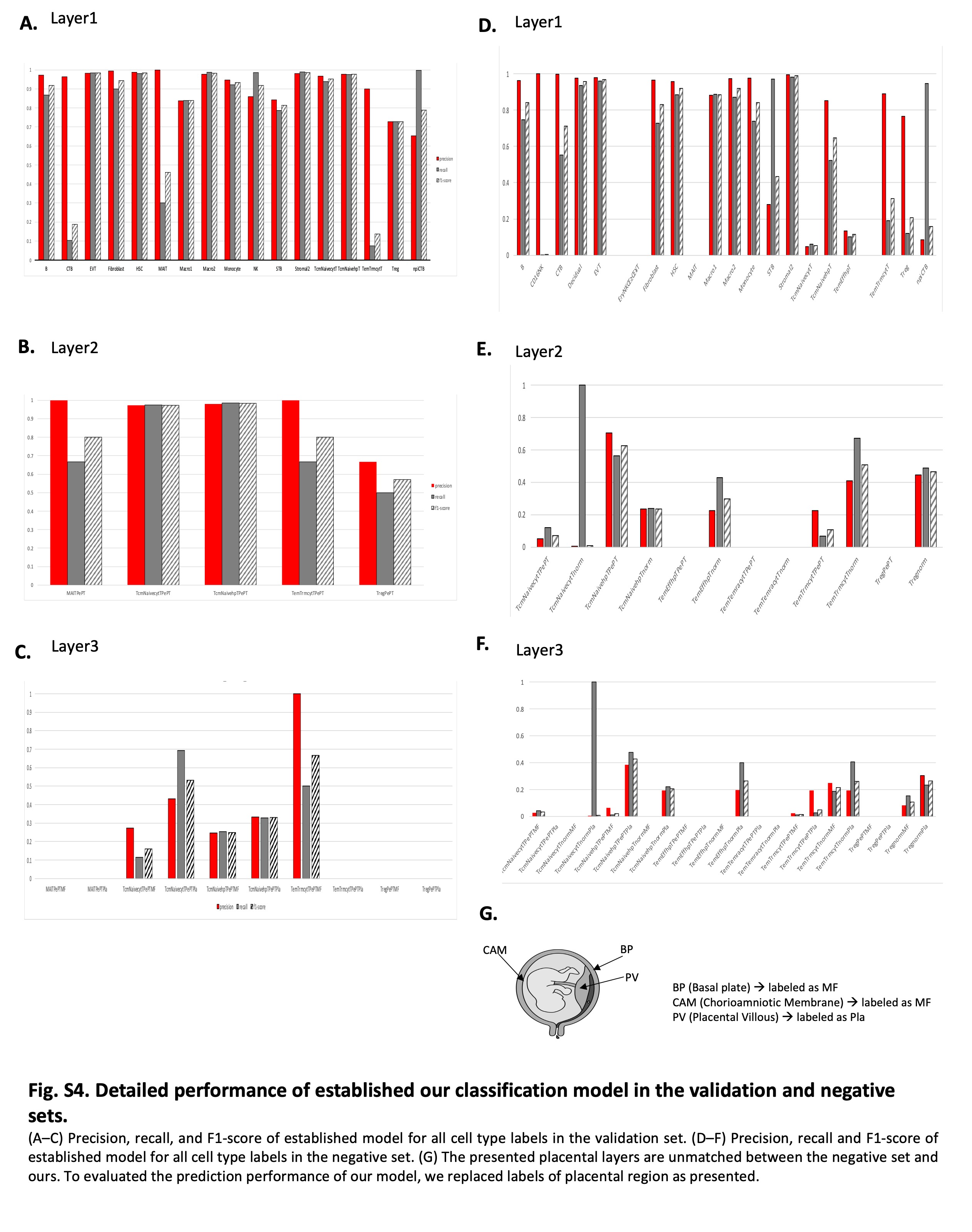

### FigS5_300.jpg

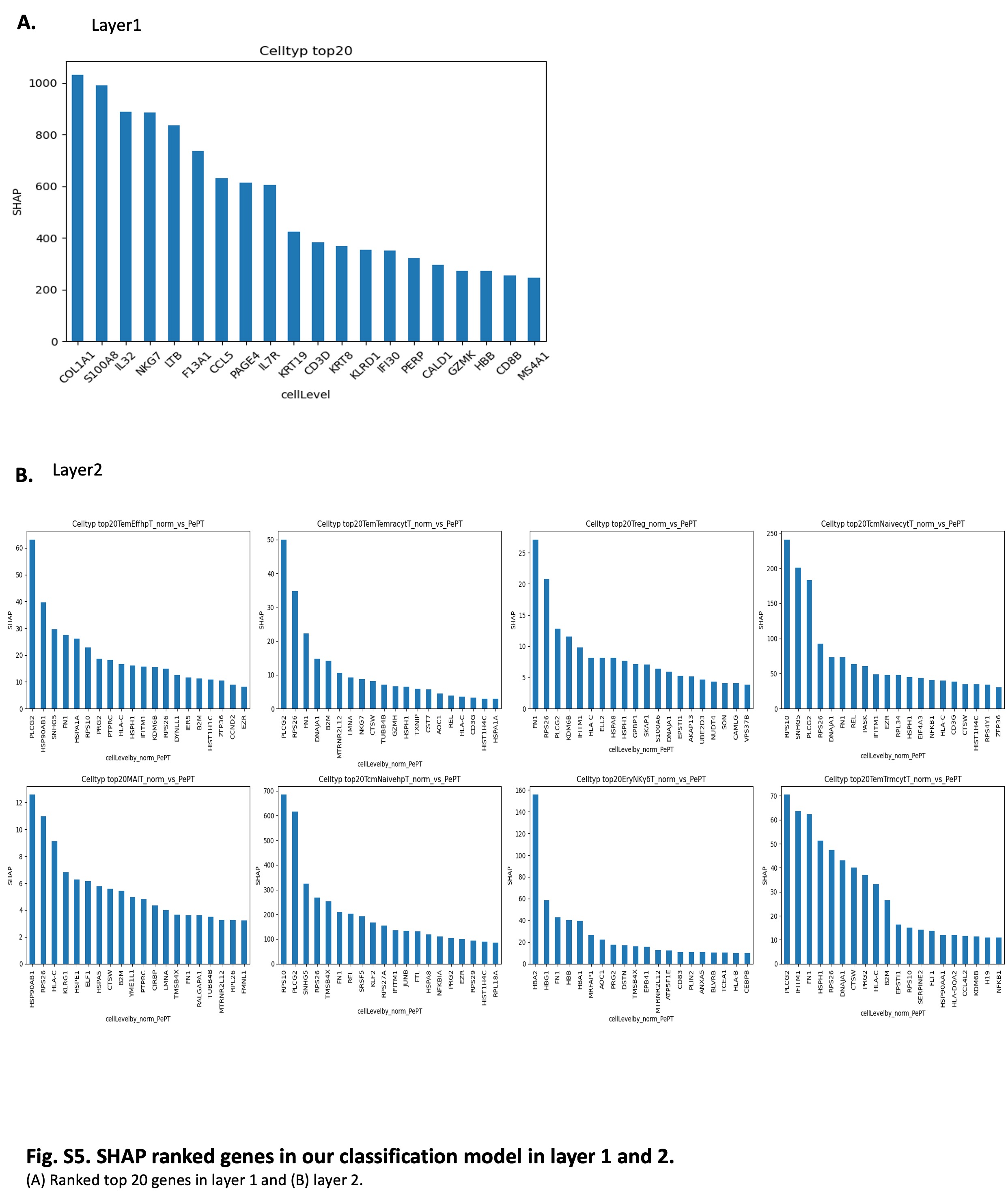

### FigS6_300.jpg

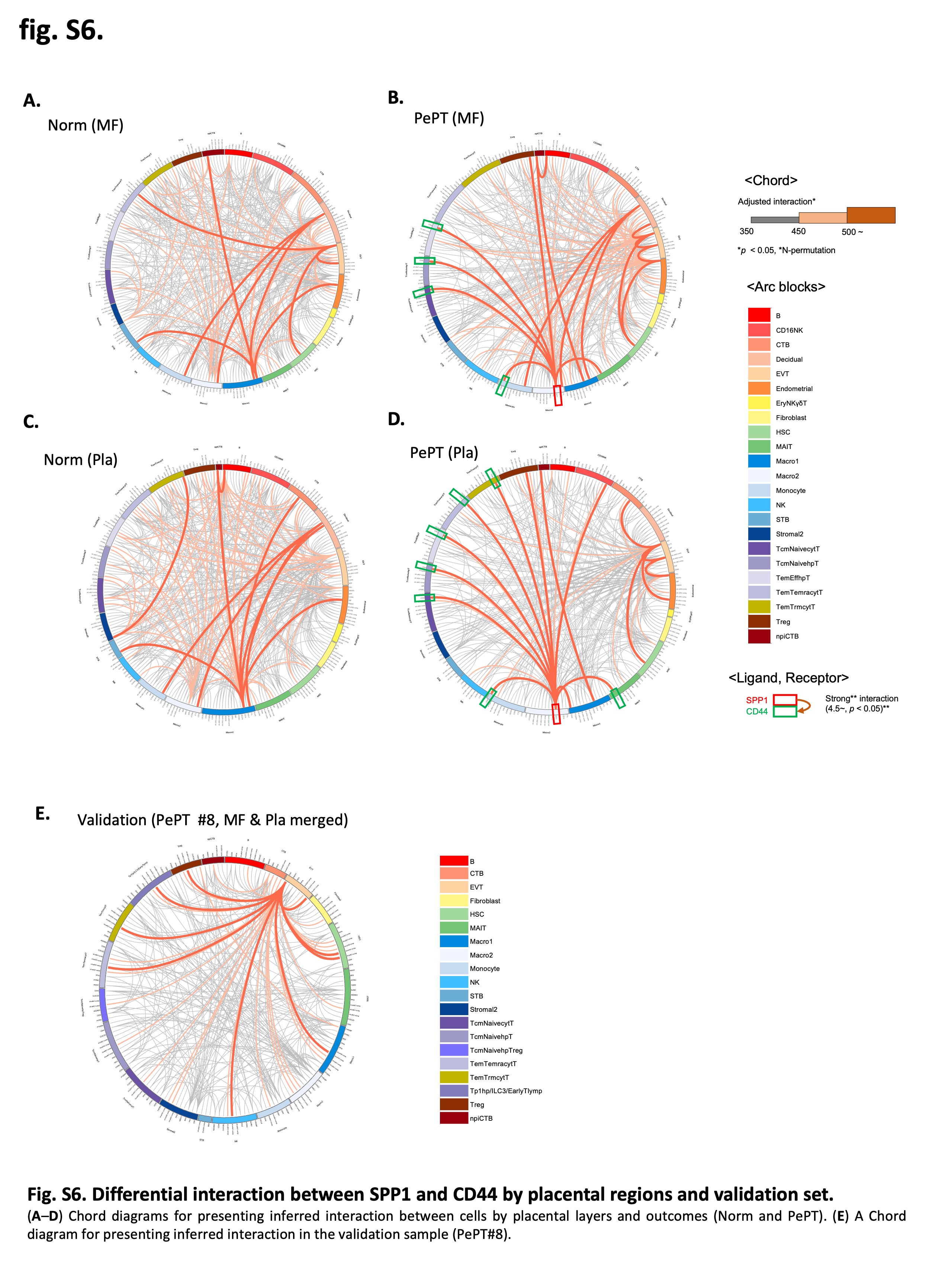

### FigS7_300.jpg

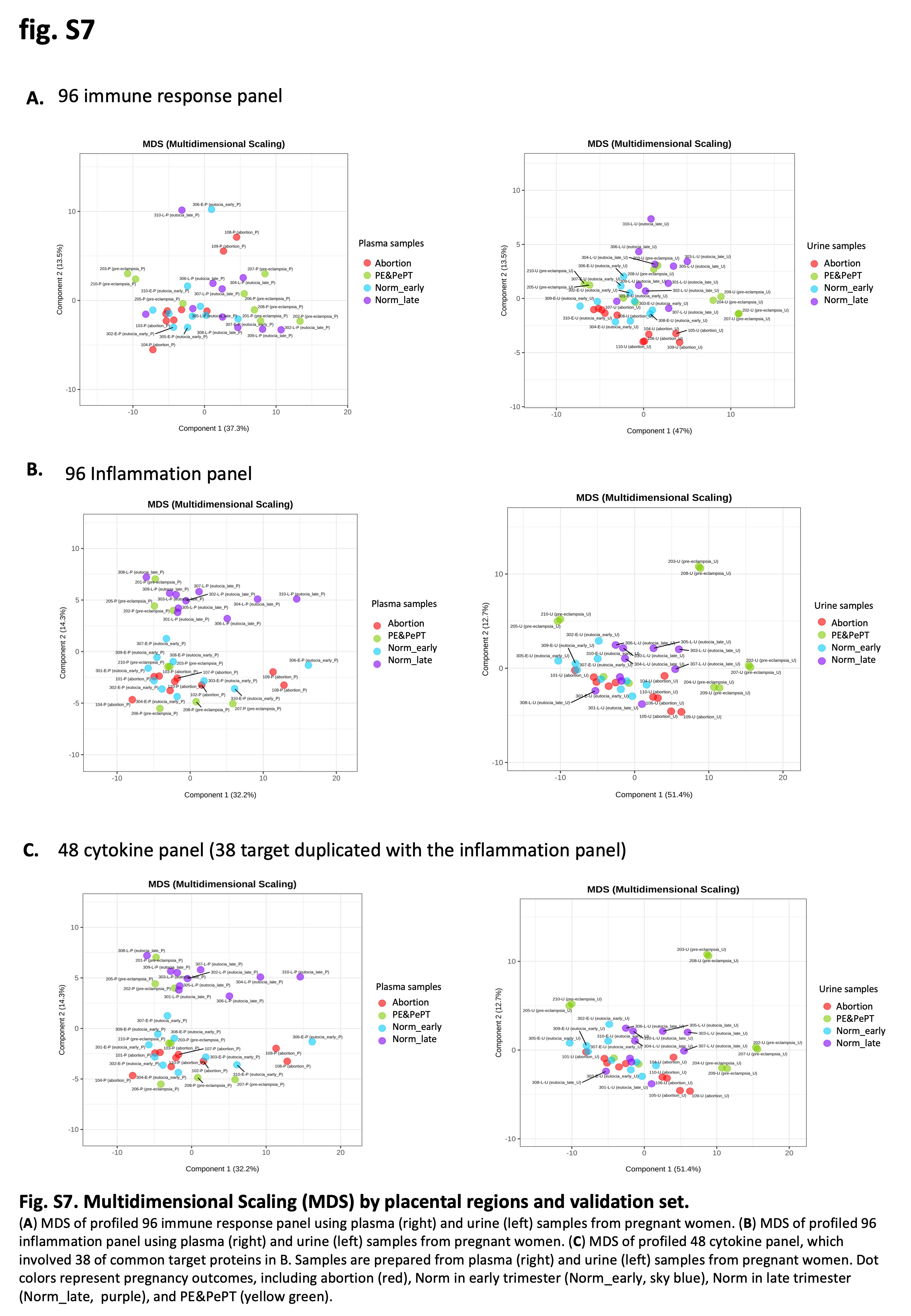
